# Hotspot of Exotic Benthic Marine Invertebrates Discovered in the Tropical East Atlantic: DNA Barcoding Insights from the Bijagós Archipelago, Guinea-Bissau

**DOI:** 10.1101/2024.07.07.602388

**Authors:** Carlos J. Moura, Peter Wirtz, Filipe T. Nhanquê, Castro Barbosa, Ester Serrão

## Abstract

**Aim:** This study aimed to explore and document putative exotic marine benthic invertebrate species in the Bijagós Archipelago, Guinea-Bissau, to enhance understanding of marine biodiversity and address the extent of marine species introductions.

**Location:** The research was conducted in the Bijagós Archipelago, a UNESCO Biosphere Reserve located in Guinea-Bissau.

**Methods:** The study involved the region’s first scuba-diving survey of marine biodiversity. DNA barcoding was employed to assist in the identification of benthic invertebrate species. Molecular phylogenetic analyses were conducted with the available DNA barcodes to ensure accurate taxonomic assignments, detect cryptic species, and investigate the phylogeography of the taxa.

**Results:** The survey resulted in the discovery of 28 new species records for the Bijagós Archipelago, including octocorals, scleractinians, hydroids, bryozoans, barnacles, and ascidians. Among these, seven species were documented for the first time in the East Atlantic: *Stragulum bicolor*, *Tubastraea tagusensis*, *Nemalecium lighti*, *Diphasia* sp., *Amathia alternata*, *A. distans*, and *Symplegma rubra*. Molecular analyses revealed pervasive cryptic diversity within species previously listed as exotic, suggesting that some, such as the hydroids *Plumularia setacea*, *Obelia geniculata*, and *Dynamena disticha*, are not exotic due to their restricted biogeographic distributions. Many other species reported as introduced present only a few genetic lineages capable of long-distance dispersal due to human activities.

**Main Conclusions:** The study highlights considerable gaps in the knowledge of West African marine biodiversity and suggests a substantial underestimation of the anthropogenic trade in exotic marine species between the Tropical East Atlantic and the Americas, and between the Indo-Pacific and West Africa. Detailed taxonomic and genomic analyses are necessary for understanding marine exotic species’ biogeography and adaptive traits. Our findings challenge current classifications of exotic species and underscore the need for improved monitoring and management to prevent the spread of non-native marine species.

## Introduction

The anthropogenic introduction of marine species outside their native ranges has been occurring for centuries, but it has been rising in recent years due to increasing human globalization. This includes more boat traffic, floating litter, aquaculture, the aquarium trade, and the construction of canals that allow the dissemination of exotic marine species (e.g. Bax et al. 2003; Geburzi and McCarthy 2018). Once established, non-native species may cause ecological, economic, and social impacts (e.g. Occhipinti-Ambrogi, 2021). Invasive species can lead locally to the decline, replacement, or extinction of native species, loss of habitat, and changes in the structure and function of ecosystems (e.g. Bax et al. 2003; Katsanevakis et al. 2014). Moreover, the economic costs associated with managing invasive species and mitigating their impacts can be substantial, affecting industries such as fisheries, tourism, and coastal infrastructure (e.g. Pimentel et al. 2005; Cuthbert et al. 2021).

The growing recognition of the threats posed by marine exotic species has prompted international efforts to manage and mitigate their impacts. Strategies include improving biosecurity measures, enhancing monitoring and early detection systems, developing rapid response and eradication plans, and ultimately encouraging the commercial use of invasive species (e.g. Davidson et al. 2017; Giakoumi et al. 2019). Effective management requires a coordinated approach that integrates science, policy, and public awareness to address the complex challenges posed by these species (Bax et al. 2003; Hulme et al. 2008).

In this study, we present and investigate the occurrence of putative exotic invertebrate species in the Bijagós Archipelago, Guinea-Bissau (Tropical West Africa), following the first scuba-diving exploration of the subtidal marine benthic biodiversity of this supposedly pristine UNESCO Biosphere Reserve of great conservation interest. The marine invertebrates from the Bijagós archipelago presented in this study were selected based on taxonomic assignments from DNA barcoding and the inclusion of their genera in the “World Register of Introduced Marine Species (WRiMS)” database (https://www.marinespecies.org/introduced/; Costello et al. 2021). We then investigated their systematics and phylogeography through phylogenetic analyses using the available barcodes of these taxa.

We report several new species records for the Bijagós Archipelago, and the country Guinea Bissau, including octocorals, scleractinians, hydroids, bryozoans, barnacles, and ascidians. Seven of these species are confirmed for the first time in the Eastern Atlantic. We identified nominal species representing complexes of cryptic taxa, some erroneously assumed to be anthropogenically introduced across large distances. We also noted many cases of exotic species in which only a few genetic lineages reveal the ability to be dispersed across long distances. This study demonstrates the utility of DNA barcoding to investigate marine invasions, and that the coastal marine biodiversity of West Africa has been considerably overlooked.

## Materials and Methods

### Sample collection

The marine invertebrate species presented here were collected manually, primarily using a knife, by C.J.M. during a scuba-diving expedition to the Bijagós Archipelago (Guinea-Bissau) from April 27 to May 11, 2023, at depths ranging from 1 to 12 meters. One exception is a hydroid species collected in the intertidal on Kere Island by P.W. Many species were photographed *in situ*. The samples were preserved in 96% ethanol and labeled with sampling data as soon as possible after collection. In a laboratory of the University of the Algarve, Portugal, the samples were sorted by morphotype and photographed using a stereomicroscope. For each distinct morphotype collected, fragments of biological tissues (approximately 1-2 cm) were isolated, re-labeled, and preserved in separate 1.5 ml Eppendorf tubes with 96% ethanol for subsequent genetic analyses.

### DNA barcoding analyses

We conducted DNA barcoding analyses on a mix of mostly marine benthic sessile invertebrate samples collected from Western Portugal, Mauritania, and Guinea-Bissau. We aimed to obtain at least three samples of each morphotype from each main geographical region. Tissue samples, approximately 3-5 mm in height, were isolated in paper tissue, relabeled with shortcodes, and placed in 0.5 ml tubes with 70-120 μl of “QuickExtract™ DNA Extraction Solution”. The genomic DNA was then extracted by transferring the tubes to a heat block at 65°C for about 15 minutes, with frequent vortexing, followed by a 2-minute incubation at 98°C.

We used the primers LCO1490 (GGTCAACAAATCATAAAGATATTGG) and HCO2198 (TAAACTTCAGGGTGACCAAAAAATCA) (Folmer et al. 1994), and SHA (ACGGAATGAACTCAAATCATGT) and SHB (TCGACTGTTTACCAAAAACATA) (Cunningham & Buss, 1993) to amplify approximately 658 and 600 base pairs of the mitochondrial genes COI and 16S, respectively. For octocorals, we used the primers COI8414 (CCAGGTAGTATGTTAGGRGA) and COIOCT (ATCATAGCATAGACCATACC) (McFadden et al. 2011) to amplify around 664 bp of the COI gene. Additionally, a pool of different primer-tag combinations designed by Srivathsan et al. (2019, 2024) was synthesized to identify PCR products through demultiplexing after DNA sequencing. For PCR amplification, we mixed 0.25–1 µl of DNA template, 0.4 µl of each primer, 6.5 µl of “Supreme NZYTaq II 2x Green Master Mix” (Nzytech, Lisbon, Portugal), and 4.5–5.25 µl of H2O. The PCR conditions were: 95 °C for 5 minutes (one cycle), followed by 34 cycles of 94 °C for 30 seconds, 46-50 °C for 40 seconds, and 72 °C for 45 seconds, with a final extension at 72 °C for 5 minutes. PCR success was verified through agarose gel electrophoresis using 2 µl of each PCR product. The PCR products were pooled based on amplification success and the scientific relevance of the genetic material, then purified collectively using Ampure beads (Beckman Coulter). We sequenced the PCR products in a single run with a MinION sequencer (©Oxford Nanopore Technologies, Oxford, United Kingdom), using the Ligation Sequencing Kit SQK-LSK114 and an R10.4 flow cell. ONTBarcoder v2.2 (Srivathsan et al. 2024) was used to demultiplex sequence reads and assemble DNA barcodes.

The taxonomic identities of the 16S and COI barcodes obtained were initially investigated using “Nucleotide Blast” searches in GenBank (https://blast.ncbi.nlm.nih.gov/) and the “Identification Engine” available in BOLD (https://v4.boldsystems.org/). Based on these results, we selected the DNA barcodes of the Bijagós’ invertebrate genera listed in “The World Register of Introduced Marine Species (WRiMS)” (https://www.marinespecies.org/introduced/).

Finally, we retrieved from GenBank (https://www.ncbi.nlm.nih.gov/nucleotide/) and BOLD (https://v4.boldsystems.org/) all COI or 16S barcodes with phylogenetic correspondence to the taxa identified from the Bijagós through DNA barcoding. The DNA barcodes generated and available online were aligned separately by marker and taxa using the MAFFT online server (https://mafft.cbrc.jp/). After trimming the alignments, we generated Maximum-Likelihood phylogenetic trees using the PHYML server (http://atgc.lirmm.fr/phyml/; Guindon et al. 2010), with 1,000 bootstrap replicates and default settings. The resulting trees were edited in ITOL V5.0 (Letunic & Bork, 2024) and INKSCAPE V1.1.

## Results and Discussion

Twenty-eight marine species belonging to genera listed in the “World Register of Introduced Marine Species” (WRiMS) (https://www.marinespecies.org/introduced/) were identified in the Bijagós archipelago, Guinea-Bissau (Fig. 1). These species were sampled at depths ranging from 0 to 12 meters and (all except one) were identified through DNA barcoding (Table 1).

**Fig. 1.**
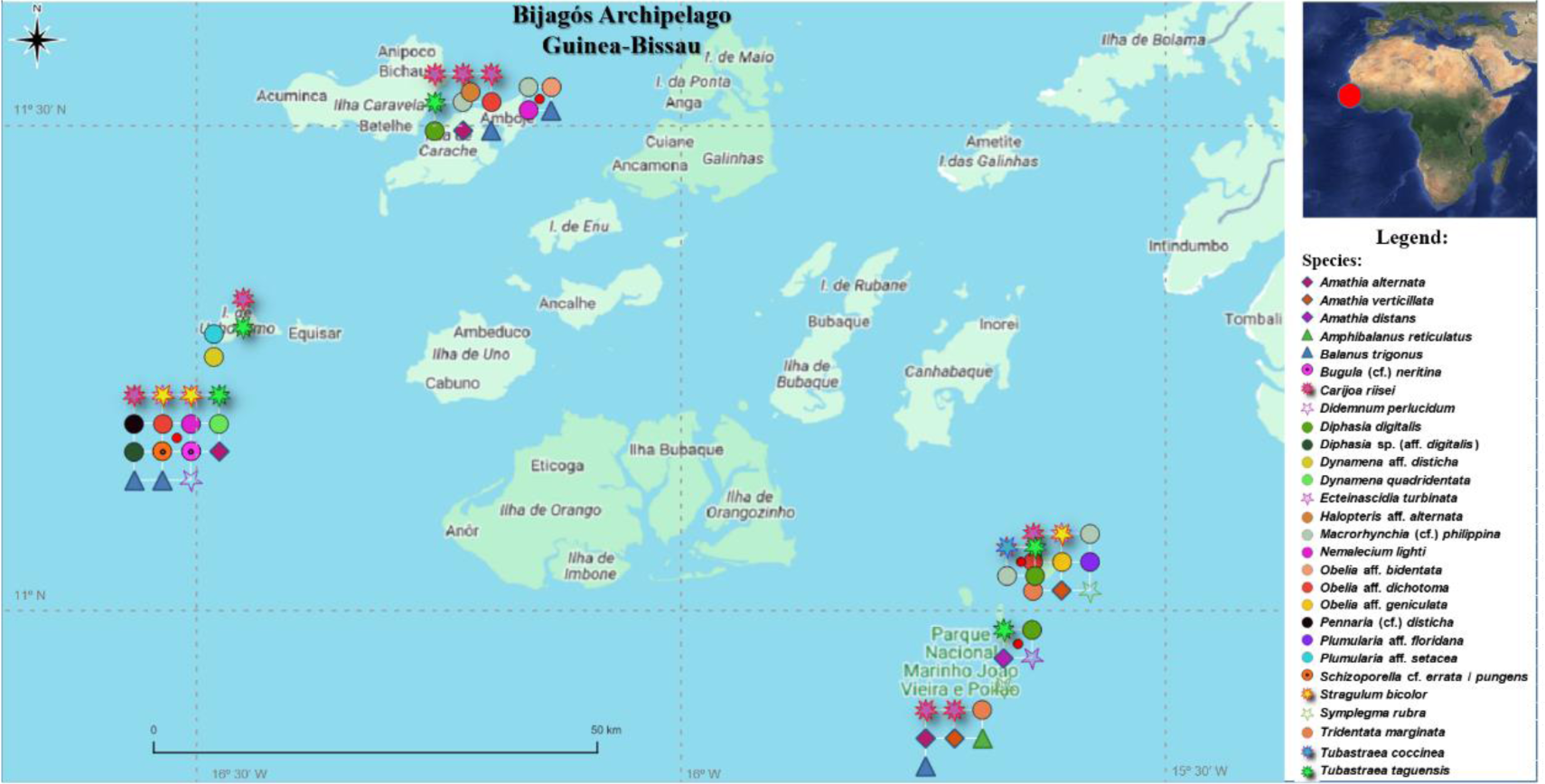
Locations in the Bijagós Archipelago (Guinea-Bissau) where the discovered species were found.

**Table 1.**
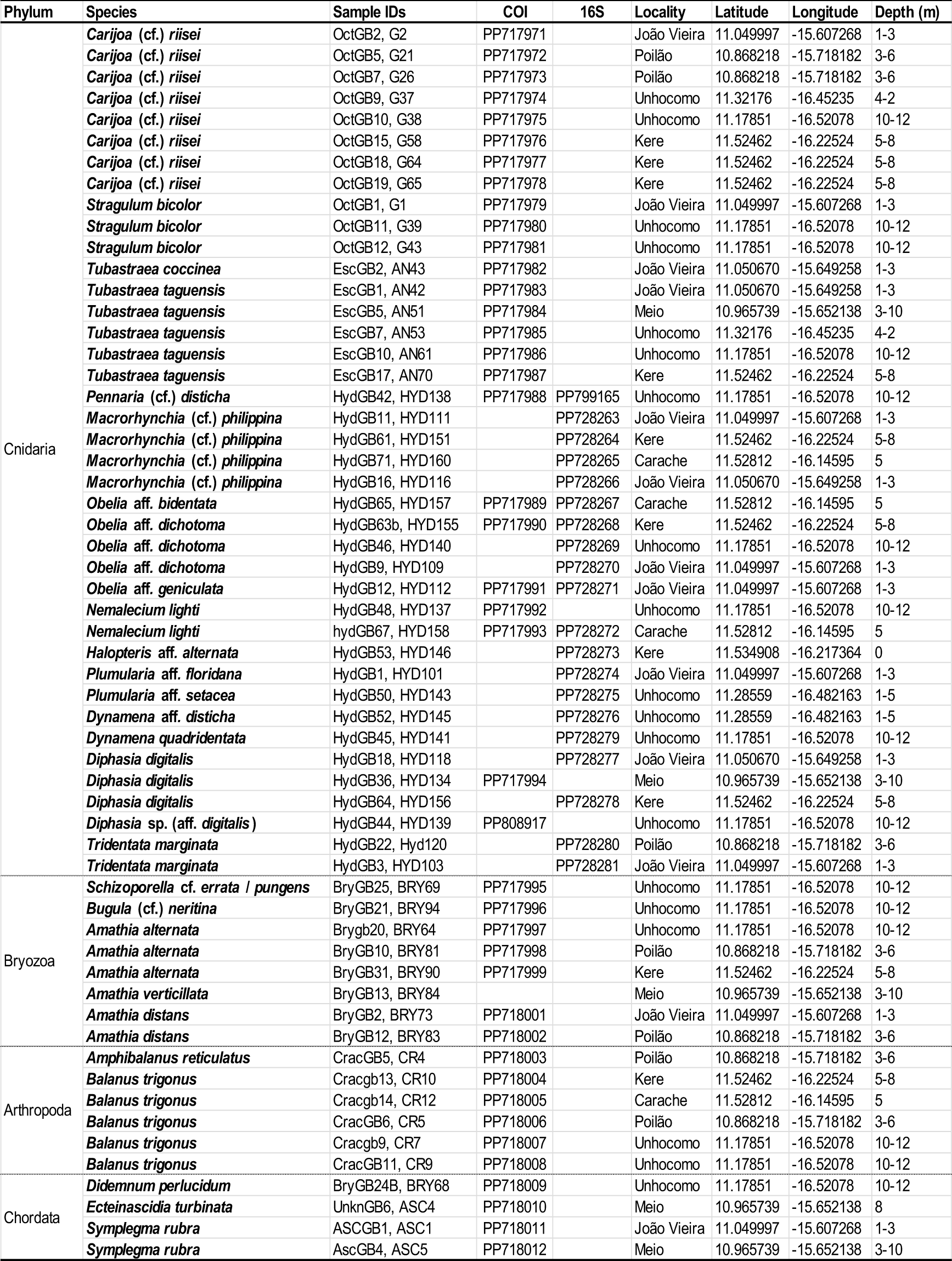
Sampling data and GenBank accession numbers of the taxa discovered in the Bijagós Archipelago.

### Taxonomic Account and Biogeography

**Phylum Cnidaria**

**Class Anthozoa**

**Order Malacalcyonacea**

**Family Carijoidae**

***Carijoa* (cf.) *riisei* (Duchassaing & Michelotti, 1860)**

(Figure 2 A-C)

**Fig. 2.**
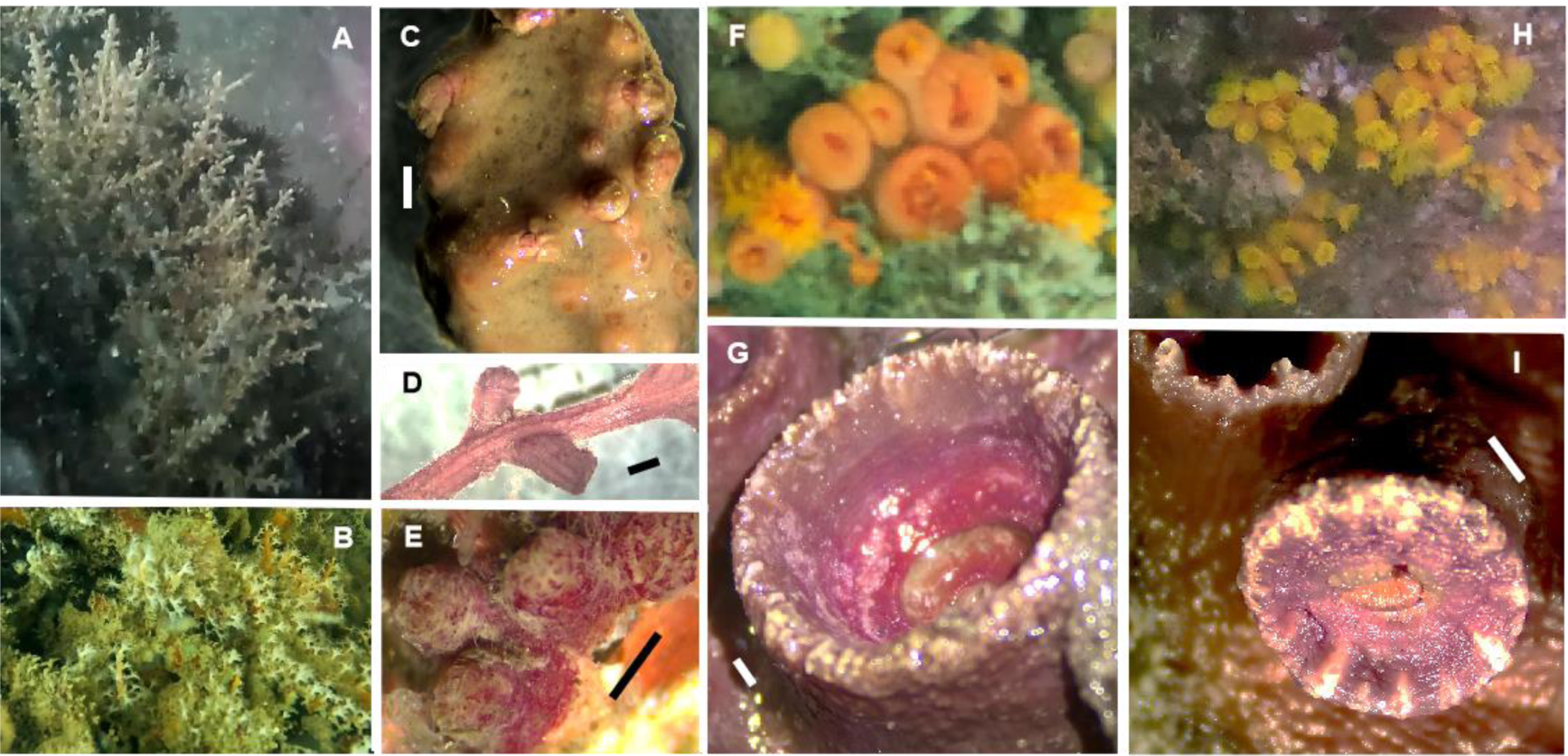
*Carijoa* (cf.) *riseii* (A-D), *Stragulum bicolor* (E), *Tubastraea coccínea* (F,G) and *T. tagusensis* (H,I). Scale bars: 1mm.

“Snowflake corals” were commonly found in the Bijagós (Table 1), either overgrown by sponges or protruding directly from rocky substrates or wrecks (Fig. 2 A-C). We sampled two COI haplotypes of *Carijoa*, differing by a single nucleotide position, found in sympatry, and not segregated by colony growth mode (Figs. S1.1). One of these COI haplotypes has been detected across tropical waters of the Pacific Ocean and in South Africa, while the other was previously found on the Pacific coast of Panama and in South Africa. Both haplotypes correspond to samples taxonomically identified as either *C. riisei* or undetermined *Carijoa* species (McFadden et al. 2012; Vargas et al. 2014; Fig. S1.1). Cryptic species may exist within the genus *Carijoa* (Concepcion et al. 2008, 2010), and given the observed differences in colony growth mode between the two COI haplotypes detected, we designate the eight octocoral colonies found in the Bijagós as “*Carijoa* (cf.) *riisei*”.

The nominal species *C. riisei*, originating from the Indo-Pacific and widely distributed across tropical and subtropical seas worldwide (Concepcion et al. 2010), has been previously reported in West Africa at São Tomé and Principe (Concepcion et al. 2010), off Gabon (Friedlander et al. 2014), Gabon (Wirtz 2020), Cape Verde Islands (Lopes et al. 2021), Senegal (pers. comm. Patrice de Voize, in Lopes et al. 2021; pers. comm. Wirtz), and Nigeria (Orrell 2024). This study marks the first record of *Carijoa* corals in Guinea-Bissau.

### Family Tubiporidae

***Stragulum bicolor* van Ofwegen & Haddad, 2011**

(Fig. 2 D)

This encrusting and inconspicuous octocoral is reported here occurring on a natural rocky substrate near João Vieira Island and overgrowing a wreck and *Tubastraea tagusensis* southwest of Unhocomo Island. The three COI barcodes generated are identical to available barcodes of *S. bicolor* from Brazil, the Caribbean, and the tropical NE Pacific (Fig. S1.2). *Stragulum* is a monotypic genus, confirming the taxonomic assignment of these three octocorals sampled from the Bijagós as *S. bicolor*.

The first taxonomic description of *Stragulum bicolor* is recent, following its initial discovery and subsequent observations on artificial substrates, suggesting its spread in Brazilian waters since the early 21^st^ century (van Ofwegen & Haddad, 2011). Subsequently, this species was recorded in the Caribbean and Persian Gulf (Samimi-Namin et al. 2022), and now in Guinea-Bissau. This marks the first report of *S. bicolor* in the East Atlantic, raising questions about its invasive potential and history as a cryptogenic species.

### Order Scleractinea

**Family Dendrophylliidae**

***Tubastraea coccinea* Lesson, 1830**

(Fig. 2 F, G)

The “orange-cup coral” was discovered near João Vieira Island, thriving on a natural rocky substrate. The COI haplotype identified in this colony matches those found in samples from Hawaii (Hellberg, 2006) and the Gulf of Mexico (Figueroa et al. 2019) (Fig. S1.3). Morphologically, the specimen collected in the Bijagós (cf. Fig. 2G) closely resembles *T. coccinea* specimens DNA barcoded from the Gulf of Mexico (Figueroa et al. 2019).

This Indo-Pacific species, widely distributed and often observed on artificial substrates, has been spreading throughout the West Atlantic since at least 1951 (Cairns, 2000). Its presence was first recorded on West African shores in 2015, specifically in the Canary Islands (Brito et al. 2017), and subsequently in São Tomé and Príncipe (Wirtz, 2023). Previous records of *T. coccinea* from the Cape Verde Islands and the Gulf of Guinea have since been reassigned to *Atlantia caboverdiana* (Ocaña & Brito, 2015) (Capel et al. 2020).

The DNA barcoding and subsequent phylogenetic analysis of this coral from the Bijagós (Fig. S1.3) confirm the first documented occurrence of *T. coccinea* in Guinea-Bissau. Further comprehensive genetic studies and monitoring efforts are necessary to elucidate the extent of its presence and potential spread along the West African Atlantic coast.

***Tubastraea tagusensis* Wells, 1982**

(Fig. 2 H, I)

The “Tagus Cove Cup Coral” was found on shipwrecks and natural rocky habitats at five locations in the Bijagós: João Vieira, Meio, Kere, and north and southwest of Unhocomo Island. These five scleractinian colonies share the same COI haplotype (Fig. S1.3), previously identified in the Gulf of Mexico as *T. tagusensis* (Figueroa et al. 2019), and in the Maldives and Yemen as *T. micranthus* (Ehrenberg, 1834). *Tubastraea micranthus* typically has a dark-green color, unlike the orange *Tubastrea* observed in the Bijagós (Fig. 2 H), which morphologically resembles the *T. tagusensis* colonies described by Figueroa et al. (2019).

This invasive coral, likely of (Indo-) Pacific origin, has been documented spreading rapidly in recent decades across both artificial and natural marine habitats in Brazil (Figueira de Paula and Creed, 2004) and the Gulf of Mexico (Figueroa et al. 2019).

To our knowledge, *T. tagusensis* was only recently reported in the East Atlantic from São Tomé and Príncipe (Wirtz, 2023), albeit based solely on underwater photographs. The present DNA barcoding analysis confirms the presence of *T. tagusensis* in the Bijagós archipelago, and thus in the Eastern Atlantic, significantly expanding its known geographical range.

### Class Hydrozoa

**Order Anthoathecata**

**Family Pennariidae**

***Pennaria* (cf.) *disticha* Goldfuss, 1820**

(Fig. 3 A)

**Fig. 3.**
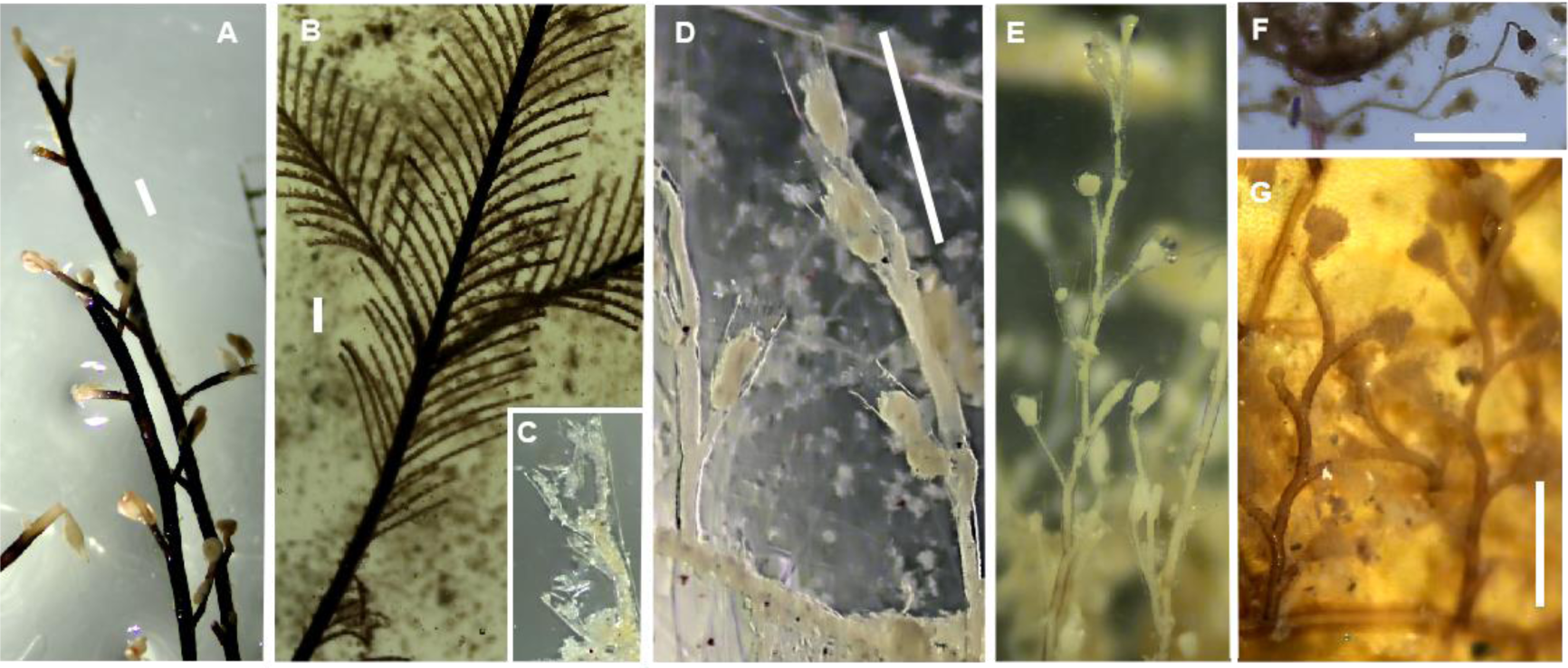
*Pennaria disticha* (A), *Macrorhynchia philippina* (B,C), *Obelia* aff. *Bidentata* (D) *Obelia* aff. *dichotoma* (E, F), *Obelia* aff. *geniculata* (G). Scale bars: 1mm.

The “Christmas tree hydroid” was discovered only once, near Unhocomo Island. Cryptic diversity likely exists within the widely distributed nominal species *P. disticha* (Miglietta et al. 2015; 2019; Vaga et al. 2020). This diversity is highlighted in our phylogenetic reconstructions, revealing two main divergent clades based on available 16S barcodes (Fig. S1.4), corresponding to at least two cryptic species, and three main evolutionary branches using COI sequence data (Fig. S1.5), corresponding to at least three distinct cryptic species. The 16S sequence of the Bijagós sample is 100% identical to the 16S barcodes of colonies from Mauritania, the Mediterranean, and the Gulf of Mexico (Fig. S1.4). These samples represent one of the more widely spread 16S haplotypes within the *P. disticha* complex, albeit restricted to the Atlantic basin (Fig. S1.4). Other 16S haplotypes within this main clade or cryptic species were detected in the Caribbean, Brazil, Azores, and Indo-Pacific (Fig. S1.4).

The availability of COI barcodes of *Pennaria* is limited (Fig. S1.5). The COI sequence of the Guinea-Bissau sample is unique and slightly distinct from a COI haplotype found in Mauritania (Fig. S1.5).

*Pennaria* species are distributed throughout warm-temperate to tropical waters worldwide (Schuchert, 2006). Previous reports in West Africa include the Cape Verde Islands (Rees & Thursfield, 1965), Madeira (Wirtz, 2007), the Canary Islands, Western Sahara, Senegal, and Gambia (GBIF.org, 2024a). This marks the first report of *Pennaria* from Guinea-Bissau.

### Order Leptothecata

**Family Aglaopheniidae**

*Macrorhynchia* (cf.) *philippina* Kirchenpauer, 1872

(Fig. 3 B, C)

The “’Stinging bush hydroid” was collected near three Bijagós islands: Karache, Kere, and João Vieira (Fig. 1). *Macrorhynchia philippina* comprises two main divergent lineages that likely correspond to two cryptic species (Moura et al. 2018): one species is restricted to the Indo-Pacific, and the other is widely dispersed across tropical to subtropical waters globally (Fig. S1.6). The four colonies collected in the Bijagós archipelago share identical 16S sequences, showing 100% similarity to specimens from Sierra Leone, Madeira, Brazil, the Caribbean, the southwestern Indian Ocean, Tahiti (South Pacific), and the Pacific Ocean coast of Panama (Fig. S1.6). The Bijagós colonies belong to the most widely dispersed 16S haplotype within the *M. philippina* complex (Fig. S1.6), likely facilitated by human-mediated transport (Moura et al. 2019). In contrast, other 16S haplotypes of this nominal species appear to have much more restricted distributions.

Along the West African coast, the “Stinging Bush Hydroid,” likely native to the Indo-Pacific (Moura et al. 2019; Fig. S1.6), has been previously reported in the Cape Verde Islands, Madeira (Ansín Agís, Ramil & Vervoort, 2001), Selvagens, Canary Islands (Riera, 2016), São Tomé and Príncipe, and Sierra Leone (Moura et al. 2018). Ansín Agís et al. (2001) state that *M. phylippina* was previously found by “Billard, 1931c, as *L. philippinus*”, in Guinea-Bissau, although we could not verify this reference. Nevertheless, the current records of *M. philippina* sensu lato (”s.l.”) from four localities confirm the species’ presence and establishment in Guinea-Bissau.

### Family Obeliidae

***Obelia* aff. *bidentata* Clark, 1875**

(Fig. 3 D)

This *Obelia* species, characterized by distinct bicuspid teeth at the hydrothecal border, was found near Carache Island, overgrowing a gorgonian and algae. According to Calder (2020), this taxon could be morphologically classified as *O. oxydentata* Stechow, 1914. However, recent molecular approaches have been inconclusive in removing *O. oxydentata* from the synonymy of *O. bidentata* (Cunha et al. 2020). While these two species do not have topotypes nominated (as suggested by Calder 2020) and are associated with DNA barcodes (as suggested by Moura et al. 2018), we cannot provide accurate taxonomic assignments.

The COI and 16S barcodes of the *Obelia* colony from the Bijagós archipelago are unique compared to available DNA barcodes of *Obelia* species. The monophyletic clade of *O. bidentata* s.l. exhibits divergent genetic lineages indicative of cryptic species diversity (Cunha et al. 2020). Phylogenetic reconstructions based on the available 16S and COI barcodes of *O. bidentata* (Figs. S1.7, S1.8) suggest the presence of four cryptic species within this complex, considering the Bijagós sample as potentially a distinct species. While the number of available DNA barcodes for this widespread species complex is limited (Figs. S1.7, S1.8), the Bijagós sample appears to represent a cryptic species with a restricted geographical distribution. Two of the four putative species identified within the *O. bidentata* complex exhibit wide biogeographic ranges, but no widespread 16S or COI haplotype has been identified within the complex (Figs. S1.7, S1.8).

The *O. bidentata* species complex is distributed circumglobally across tropical and temperate waters (Peña Cantero and García Carrascosa, 2002) and has been reported along the Atlantic coast of Africa, including Morocco (Patriti, 1970), the Canary and Cape Verde Islands (Medel & Vervoort, 2000), South Africa (Millard, 1975), Angola, Ghana, and offshore of Guinea-Bissau (GBIF, 2024b). This report of *Obelia* aff. *bidentata* represents the first record for Guinea-Bissau.

***Obelia* aff. *dichotoma* (Linnaeus, 1758)**

(Figs. 3 E, F)

Three *Obelia* specimens, morphologically resembling *Obelia dichotoma*, were discovered near Kere, João Vieira, and Unhocomo Islands. Analysis of their 16S and COI sequences suggests that colonies from Kere and João Vieira Islands belong to a distinct *Obelia* species, clustering with an *Obelia* specimen from Mauritania (Figs. S1.7, S1.8). The *Obelia* specimen from Unhocomo Island belongs to a different cryptic species, clustering with haplotypes potentially from another species found in the Gulf of Mexico and Mauritania (Figs. S1.7, S1.8).

Within this species complex, three species are identified with wide geographical distributions that suggest human-mediated transport: one found in the Gulf of Mexico, Brazil, and China; another present in Brazil, eastern Canada, Italy, and the U.K.; and a third detected in Brazil and eastern Canada (Figs. S1.7, S1.8). The two cryptic species of *Obelia* reported from the Bijagós, morphologically similar to *O. dichotoma*, appear to be restricted to the Tropical East Atlantic and thus cannot be classified as exotic species.

*Obelia dichotoma* s.l. has frequently been reported along the West African Atlantic coasts (cf. Medel and Vervoort, 2000), including offshore of the Bijagós (GBIF 2024c). Nevertheless, this study provides the first evidence of two cryptic species with morphological similarities to *O. dichotoma* from the West African coast.

***Obelia* aff. *geniculata* (Linnaeus, 1758)**

(Fig. 3 G)

This *Obelia* species resembling *Obelia geniculata*, was discovered near João Vieira Island. The hydroid colony exhibits distinct 16S and COI sequences compared to the available DNA barcodes of *Obelia* in public databases (Figs. S1.7, S1.8). It belongs to a (pseudo-)cryptic *Obelia* species (Cunha et al. 2017), likely undescribed or currently synonymized, which includes barcodes from Brazil and the Mediterranean (Figs. S1.7, S1.8). Despite its extensive amphi-Atlantic geographical distribution, no shared haplotypes have been identified across these marine regions, suggesting that this putative *Obelia* species occupies a natural biogeographical range. Based on the currently available DNA barcodes, there is no indication of human-mediated transport for any species within the *Obelia geniculata* complex (Figs. S1.7, S1.8).

*Obelia geniculata* s.l. has been commonly observed along the West African shores, including Morocco, Madeira, Senegal, the Gulf of Guinea, Congo, and South Africa (cf. Medel & Vervoort, 2000). Nonetheless, this represents the first record of this nominal species in Guinea-Bissau.

### Family Haleciidae

***Nemalecium lighti* (Hargitt, 1924)**

(Fig. 4 A)

**Fig. 4.**
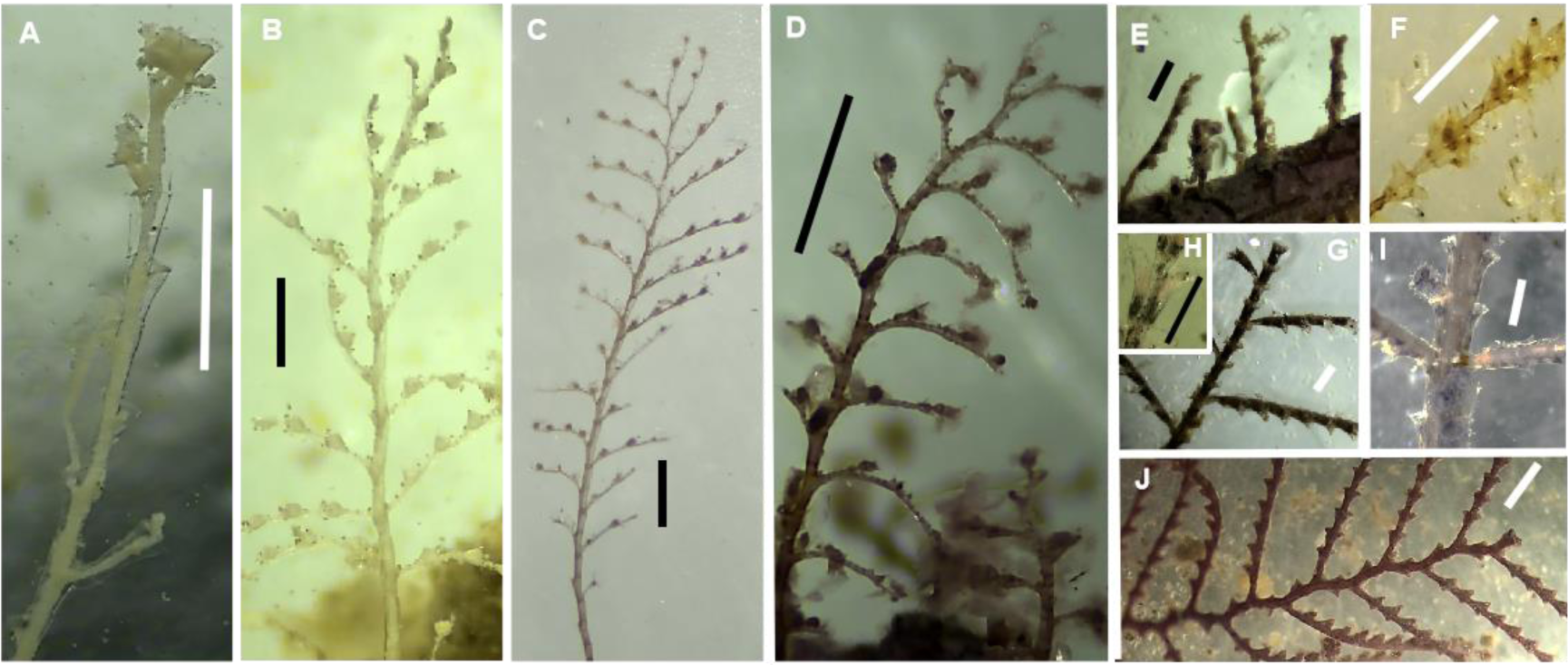
*Nemalecium lighti* (A), *Halopteris* aff*. alternata* (B), *Plumularia* aff*. Floridana* (C) *Plumularia* aff*. setacea* (D), *Dynamena disticha* (E), *Dynamena quadridentata* (F), *Diphasia digitalis* (G,H), *Diphasia* sp. (aff. *digitalis*) (I), *Sertularia marginata* (J). Scale bars: 1mm.

This species was discovered in the vicinity of Karache and Unhocomo Islands. The two hydroid colonies collected exhibit COI sequences identical to those of Caribbean colonies documented by Macher et al. (2021); Fig. S1.9). Interestingly, specimens from the Canary Islands and the Cape Verde Islands cluster in a separate clade of the same species with a COI barcode matching a sample from Hawaii, indicating multiple introductions of *N. lighti* into the Tropical East Atlantic from the Indo-Pacific (Fig. S1.9).

The 16S sequence identified in the Karache specimen represents a unique barcode, nearly identical to the 16S barcodes of Caribbean specimens (Fig. S1.10). Cryptic diversity has been observed within the genus *Nemalecium* (Boisson et al. 2016). The phylogenetic tree constructed with all available 16S barcodes of *N. lighti* reveals three distinct divergent clades, indicating at least three cryptic species (Fig. S1.10). The *Nemalecium* colonies from the Bijagós belong to the more widely distributed cryptic species within the *N. lighti* complex, found across warm waters of the Indo-Pacific, West Atlantic, and West Africa (Fig. S1.10), suggesting long-distance dispersal facilitated by rafting on boat hulls. The other putative cryptic species appear to have much more restricted biogeographical distributions (Fig. S1.10).

This study presents the first reports of the Indo-Pacific nominal species *N. lighti* in the East Atlantic, indicating a relatively recent invasion of the African Atlantic shores. However, it is noteworthy that the first record of this species in the West Atlantic dates back to 1983 in Bermuda (Calder, 1991). The current phylogenetic analyses suggest multiple invasions of this species into the East Atlantic, and while it may have been overlooked along African shores for a considerable period, we cannot dismiss the possibility of a trans-Atlantic invasion directed eastwards in this case.

### Family Halopterididae

***Halopteris aff. alternata* (Nutting, 1900)**

(Fig. 4 B)

One *Halopteris* species morphologically similar to *H. alternata,* was discovered on rocks in the intertidal zone of Kere Island. Cryptic diversity exists within the nominal species *H. alternata* s.l. (Moura et al. 2018). The 16S barcode of the Bijagós colony corresponds to “*Halopteris alternata* - lineage 5” (*sensu* Moura et al. 2018) (Fig. S1.11), which represents the most widely distributed cryptic species within the *H. alternata* complex (Moura et al. 2018). The 16S sequence from the Bijagós is identical to sequences previously documented from NW African islands (Madeira, Selvagens, São Tomé and Príncipe), Brazil, the Caribbean, and the Tropical East Pacific (Fig. S1.11). Although this lineage has not been observed in ports or marinas, its rafting capabilities, possibly facilitated by boat hulls, likely account for its recent geographical expansion (Moura et al. 2019).

The nominal species *H. alternata* has been reported previously along the West African coasts, including Madeira, Canary Islands, Cape Verde Islands (Ansín Agís et al. 2001), Selvagens, and São Tomé and Príncipe (Moura et al. 2018). This represents the first documented occurrence of this taxon in Guinea-Bissau.

### Family Plumulariidae

***Plumularia* aff. *floridana***

(Fig. 4 C)

This species, bearing strong morphological resemblances to *P. floridana* and *P. setacea*, was collected near João Vieira Island. The sample exhibits a distinct 16S haplotype that clusters within the *P. floridana* species complex (cf. Moura et al. 2018; Fig. S1.12). The unique 16S haplotype found in the Bijagós sample likely indicates the presence of one cryptic species within its naturally restricted biogeographical range in the Tropical East Atlantic (Fig. S1.12).

The nominal species *P. floridana*, purportedly distributed globally, has previously been reported in West Africa only in the Canary Islands and Cameroon (Ansín Agís et al. 2001). This marks the first documented occurrence of this species in Guinea-Bissau. Many previous reports of *P. floridana* s.l. in the literature may have been misclassified under the nominal species *P. setacea*.

***Plumularia* aff. *setacea* (Linnaeus, 1758)**

(Fig. 4 D)

This *Plumularia* species, morphologically resembling *P. setacea*, was collected on the north coast of Unhocomo Island. *Plumularia setacea* comprises a cryptic species complex (Schuchert, 2014; Moura et al. 2018). The colony from Bijagós exhibits a unique 16S haplotype, previously unreported, indicating a new cryptic species of *Plumularia* (Fig. S1.12). It clusters within a clade with samples identified as *P. setacea*, *P. strictocarpa*, *P. warreni*, *P. duseni*, *P. virginiae*, and *P. lagenifera* (Fig. S1.12). This cryptic species from the Bijagós appears to be within its naturally restricted biogeographical range.

The purportedly cosmopolitan nominal species *P. setacea* has been frequently detected along the Atlantic shores of West Africa, including Guinea-Bissau (cf. Ansín Agís et al. 2001). However, the cryptic *Plumularia* identified in Bijagós differs from four other cryptic “*P. setacea*” species previously DNA barcoded from other West African islands (Madeira, Selvagens, Tenerife, and São Tomé and Príncipe) (Fig. S1.12).

Analysis of the available 16S barcodes of the *P. setacea* complex does not suggest long-distance dispersal of *Plumularia* species (Fig. S1.12). While some colonies have been observed overgrowing artificial structures in ports elsewhere (e.g., Moura et al. 2018), the considerable genetic lineage or species segregation by geographical area (Fig. S1.12) suggests that species within the *P. setacea* complex are sensitive to substantial environmental changes, potentially limiting their dispersal ability.

### Family Sertulariidae

***Dynamena* aff. *disticha* (Bosc, 1802)**

(Fig. 4 E)

This species was detected on the north shore of Unhocomo Island, thriving on sargassum. Its 16S haplotype, previously unreported, falls within the cryptic species complex of *D. disticha* (cf. Moura et al. 2011) and shows closer phylogenetic affinity to a lineage represented by a colony from the Caribbean (Fig. S1.13). Phylogenetic analysis using the limited available 16S barcodes of *D. disticha* s.l. (Fig. S1.13) reveals lineage segregation by geographical area and does not suggest successful human-mediated dispersal over long distances.

The nominal species *D. disticha* has been documented along the West African shores, including Morocco, Madeira, Western Sahara, Mauritania, Cape Verde Islands, Senegal, Guinea-Bissau, Ghana, and Guinea (cf. Gil & Ramil 2023, and Moura et al. 2011). However, further DNA barcoding is essential to differentiate species or genetic lineages within the *D. disticha* complex reported in these regions.

***Dynamena quadridentata* (Ellis & Solander, 1786)**

(Fig. 4 F)

This taxon was discovered on a wreck southwest of Unhocomo Island. Its 16S sequence matches those of *D. quadridentata* sampled in Mauritania and the Azores (Fig. S1.13), indicating probable long-distance dispersal facilitated by rafting on algae and potentially boat hulls.

*Dynamena quadridentata* has a circumglobal distribution across temperate and tropical waters (Vervoort & Watson 2003), and in Tropical West Africa was previously reported only in Ghana (cf. Gil & Ramil, 2023). Therefore, this report marks the first record of *D. quadridentata* in Guinea-Bissau.

***Diphasia digitalis* (Busk, 1852)**

(Fig. 4 G, H)

This species was found near João Vieira, Kere, and Meio Islands. The 16S barcodes generated for specimens from João Vieira and Kere are identical and differ by three nucleotide positions from a lineage represented by four 16S barcodes from Mauritania (Fig. S1.14). The COI barcode of the colony from Meio matches five COI barcodes of specimens from Mauritania (Fig. S1.15).

*Diphasia digitalis* is a circumglobal species, historically documented along the West African Atlantic coasts of Guinea-Bissau, Guinea, Ivory Coast, and Gabon (Gil & Ramil 2017, 2023). Despite old records from West Africa, its recent discovery in the Mediterranean flagged this species as exotic (Morri et al. 2009).

***Diphasia* sp. (aff. *digitalis*)**

(Fig. 4 I)

Distinct yet morphologically similar to *D. digitalis*, this species exhibits different gonothecae (reproductive structures). It was discovered on a wreck southwest of Unhocomo Island. Its COI barcode shows eleven nucleotide differences compared to the available COI barcode of *D. digitalis* from Guinea-Bissau (Fig. S1.15). This finding highlights the probable native occurrence of this species, alongside *D. digitalis*, in the tropical waters of West Africa.

***Tridentata marginata* (Kirchenpauer, 1864)**

(Fig. 4 J)

This species was discovered near João Vieira and Poilão Islands, spreading over rocky substrates. The 16S barcodes of these colonies are identical but distinct from currently available 16S barcodes of *T. marginata* samples from Mauritania, Brazil, the Azores, and Madeira (Fig. S1.16). More DNA barcodes of this species across its distribution range are needed to enhance understanding of its potential biological invasions through phylogeographic analysis.

*Tridentata marginata* exhibits a circumglobal distribution in tropical and subtropical seas (e.g. Medel & Vervoort 1998) and has been frequently observed along the West African coastline, including Morocco, Mauritania, Cape Verde Islands, Guinea-Bissau, Guinea, Ghana, and Congo (Medel & Vervoort 1998; Gil & Ramil 2017, 2023).

### Phylum Bryozoa

**Class Gymnolaemata**

**Order Cheilostomatida**

**Family Schizoporellidae**

***Schizoporella* cf. *errata* (Waters, 1878) / *pungens* Canu & Bassler, 1928**

(Fig. 5 A)

**Fig. 5.**
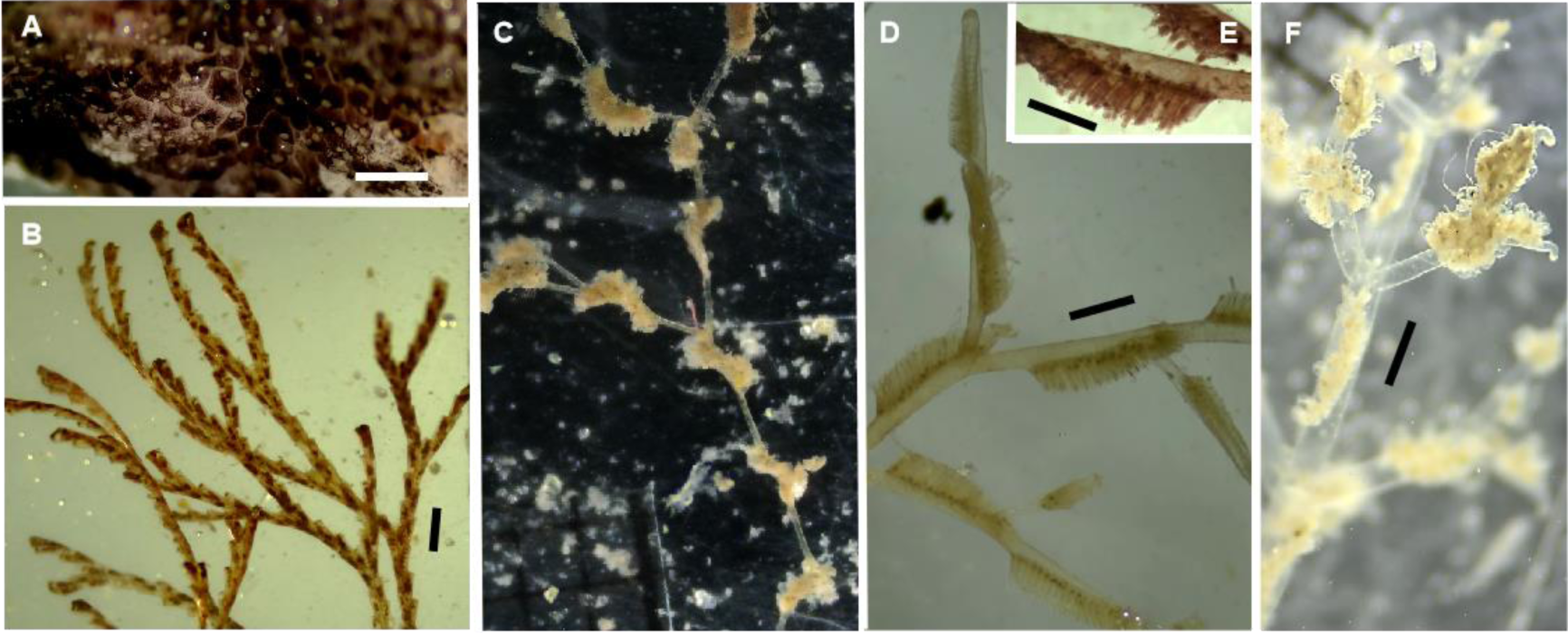
*Schizoporella pungens* (A), *Bugula* (cf.) *neritica* (B), *Amathia* (cf.) *distans* (C), *Amathia* (cf.) *alternata* (D, E), and *Amathia* sp. (F). Scale bars: 1mm.

A single colony of *Schizoporella* was found on a wreck at a depth of 10-12 meters southwest of Unhocomo Island. Its COI barcode matches those of samples identified as either *S. errata* or *S. pungens* from the Pacific, Mediterranean, and Mauritania (Fig. S1.17). This COI lineage appears to be the most widely distributed and potentially invasive among all *Schizoporella* species (Fig. S1.17), likely facilitated by rafting on boat hulls. Phylogenetic analysis incorporating all available COI barcodes of this genus (Fig. S1.17) suggests that the nominal species *S. errata* and *S. pungens*, which are morphologically similar (e.g. McCan et al. 2019), may be synonymous, with *S. errata* having nomenclatural priority. The dubious morphologic differentiation between the nominal species *S. pungens* and *S. errata*, and also *S. pseudoerrata*, *S. erratoidea*, *S. mazatlantica*, and *S. variabilis* (plus *S. serialis* and *S. isabelleana* in the synonymy of *S. pungens*, and *S. violacea* in the synonymy of *S. errata*) (McCan et al. 2019), implies that *Schizoporella* species of that complex need taxonomic reassessment, and are better identified through DNA barcoding.

Either *S. errata* or *S. pungens* (*sensu lato*) are widespread species across tropical and subtropical marine waters worldwide, often reported in ports or marinas (e.g. Marques et al. 2013; Canu & Bassler, 1930). Reports of *S. errata* along the West African coasts date from 1946 in Angola (Pestana et al. 2017), followed by reports from Ghana (in 1970; Taylor et al. 2008) and the Canary Islands (Arístegui & Cruz, 1986). The only report of *S. pungens* on the West African coasts, from 2010, is from Madeira (Canning-Clode et al. 2013). According to the phylogeny with the available COI barcodes of the genus *Schizoporella* (Fig. S1.17), the specimens previously identified from the Atlantic basin (including the Mediterranean) either as *S. pungens* or *S. errata*, most likely correspond to the same species. This is the first record of this taxon in Guinea-Bissau, which may be much more widespread in the East Atlantic than assumed.

### Family Bugulidae

***Bugula* (cf.) *neritina* (Linnaeus, 1758) (sensu latu)**

(Fig. 5 B)

The nominal species *B. neritina* was collected only once at the Bijagós, on a wreck southwest of Unhocomo Island. Cryptic species exist with3in the *B. neritica* complex (Mcgovern & Hellberg 2003). The COI barcode determined for this sample belongs to the most widespread lineage of the nominal species - the “Southern / shallow clade” (*sensu* Mcgovern & Hellberg 2003) (Fig. S1.18). The lineage named “Deep clade” seems restricted to the NE Pacific, and the lineage named “Northern clade” occurs across the North Atlantic, and in Australia (Mcgovern & Hellberg 2003; Fehlauer-Ale et al. 2014; Fig. S1.18). The COI haplotype detected in Guinea-Bissau is the most widespread, also occurring in Mauritania, South Africa, Brazil, NE Pacific, Australia, New Zealand, the Mediterranean, W Portugal, NW Spain, and the UK (Fig. S1.18). Such widespread distribution of a bryozoan COI haplotype indicates this species has been transported rottenly on boat hulls across cold to tropical waters.

*Bugula neritina*, originally described from the Mediterranean by Linnaeus in 1758, has been documented worldwide in temperate to tropical marine areas, and has long been observed along the West African coast, including Senegal (in the 1840s, US National Museum of Natural History sample 5595), Madeira (Norman 1909; Ramalhosa et al. 2017), the Canary Islands (Moro et al. 2003), Angola (e,g, Pestana et al. 2020), and Cape Verde Islands (Castro et al. 2023), This study represents the first report of *B. neritina* from Guinea-Bissau.

### Order Ctenostomatida

**Family Vesicularioidea**

***Amathia alternata* Lamouroux, 1816**

(Fig. 5 D, E)

This bryozoan species was collected near Unhocomo, Poilão, and Kere Islands. The specimens from these locations share identical COI barcodes with specimens from Brazil (Genbank accession: OK120385; Nascimento et al. 2022) and Mauritania (Fig. S1.19). However, COI barcodes from two specimens collected in Virginia, USA (Genbank accessions: OQ322772, OQ322979; Aguilar et al. direct submission), were recently classified as *A. alternata*. These two COI barcodes cluster with specimens from Brazil classified as *A. brasiliensis* (Fehlauer-Ale et al. 2011) (Fig. S1.19), which suggests misidentified samples because Fehlauer-Ale et al. (2011) classified *A. brasiliensis* and *A. alternata* from Brazil based on type-specimen comparisons.

*Amathia alternata* has been documented exclusively in the West Atlantic, ranging from North Carolina to the Caribbean, the Gulf of Mexico, and Brazil more recently (cf. Nascimento et al. 2022). This study marks the first report of *A. alternata* in the East Atlantic.

***Amathia distans* Busk, 1886**

(Fig. 5 C)

This species was collected near João Vieira and Poilão Islands. At João Vieira, it was observed infesting an extensive intertidal rocky area, while the specimen from Poilão was found overgrowing a bivalve. The COI barcodes of the two samples of *A. distans* from the Bijagós, along with a sample identified as *A. distans* from Brazil (Genbank accession: OR632417; Decker et al. 2024), share identical sequences (Fig. S1.19). This haplotype is closely clustered with a specimen from Brazil (Genbank accession: JF490058; Fehlauer-Ale et al. 2011) classified as *A. distans* and differing in eight nucleotide positions (Fig. S1.19).

According to Fehlauer-Ale et al. (2011), “*Amathia distans* sensu stricto occurs at least from Brazil to Florida” and has doubtful reports from Australia, Indonesia, Southern California, the Gulf of California, and New Zealand. However, the website https://invasions.si.edu/ contradicts these statements, citing old reports of *A. distans* in West Africa, specifically Senegal (1973, d’Hondt 1975) and Sierra Leone (1946, Cook 1968), as well as in NW Spain, the Western Mediterranean, the Red Sea, and the Indo-Pacific. This study confirms the presence of *A. distans* in the East Atlantic.

***Amathia verticillata* (delle Chiaje, 1822)**

(Fig. 5 F)

The “spaghetti bryozoan” was collected once near Meio Island, southwest of the Bijagós. No COI barcode was obtained for this taxon; however, recent global genetic analyses of this species have indicated no signs of cryptic diversity (Nascimento et al. 2021). *Amathia verticillata* presents only one haplotype that is widely distributed worldwide, likely under the influence of boat traffic. The remaining genetic lineages apparent restrict regional distributions (Nascimento et al. 2021).

Previous reports of this cosmopolitan invasive species along the West African coast include sightings in Madeira (Wirtz and Canning-Clode, 2009), the Canary Islands (Minchin, 2012), Cape Verde Islands (Waters, 1916), Ghana (Cook, 1985), Angola (Pestana et al. 2020), and Senegal (Wirtz, 2021). This study represents the first record of *A. verticillata* from Guinea-Bissau.

### Phylum Arthropoda

**Class Maxillopoda-Order**

**Balanomorpha**

**Family Balanidae**

***Amphibalanus reticulatus* (Utinomi, 1967)**

(Fig. 6 A)

**Fig. 6.**
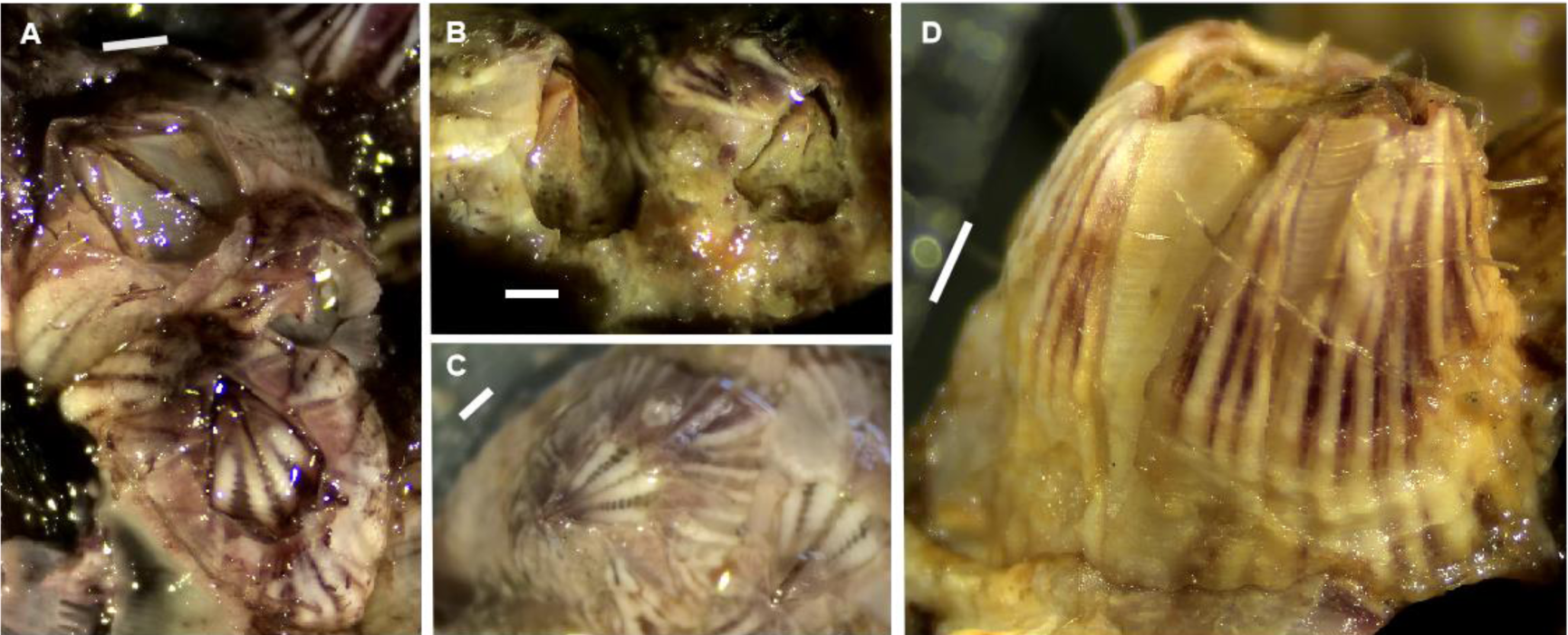
*Amphibalanus reticulatus* (A), *Balanus trigonus* (B, C, D). Scale bars: 1mm.

The “reticulated barnacle” was discovered near Poilão Island in a single occurrence. The COI barcode of the Bijagós sample shows a 100% similarity to sequences of *A. reticulatus* from Indonesia, Alaska, and Guam (Fig. S1.20), indicating it belongs to the most widely distributed lineage of this species identified through DNA barcoding. The persistence and adaptability of this particular *A. reticulatus* lineage across both cold and tropical waters are noteworthy, likely facilitated by transportation on ship hulls.

The Indo-Pacific species *A. reticulatus* has been documented in the Atlantic Ocean since the mid-20^th^ century (Henry and McLaughlin, 1975). Previous records of this invasive barnacle along the West African coast include sightings in Nigeria, Sierra Leone (Henry and McLaughlin, 1975), The Gambia (Kerckhof et al. 2010), and South Africa (GBIF, 2024d). This study marks the first documentation of *A. reticulatus* (*senso lato*) in Guinea-Bissau.

***Balanus trigonus* Darwin, 1854**

(Fig. 6 B, C, D)

The “triangle barnacle” was identified at four locations in the Bijagós: near Kere, Carache, Poilão, and Unhocomo Islands. Specimens in the Bijagós were observed on natural rocky substrates, except those near Unhocomo, which were found on a wreck and overgrowing the hydroid *Pennaria disticha*. Interestingly, all five COI sequences obtained from Guinea-Bissau samples are distinct not only from each other but also from the 24 COI barcodes of *B. trigonus* available, which represent specimens from the Pacific Ocean, NW Atlantic, and Spain (Fig. S1.21). This suggests multiple introductions of *Balanus trigonus* from the Indo-Pacific into the Tropical East Atlantic. *Balanus trigonus* is frequently documented along the West African Atlantic coasts, including Madeira, Guinea, Sierra Leone, Cameroon, Congo (Zullo, 1992), Angola (Pestana et al. 2017), Canary Islands (Gonzalez et al. 2011), Mauritania, and South Africa (https://invasions.si.edu/). This report marks the first documentation of the exotic species *B. trigonus* in Guinea-Bissau.

### Phylum Chordata

**Class Ascidiacea**

**Order Aplousobranchia**

**Family Didemnidae**

***Didemnum perlucidum* Monniot F., 1983**

(Fig. 7 A)

**Fig. 7.**
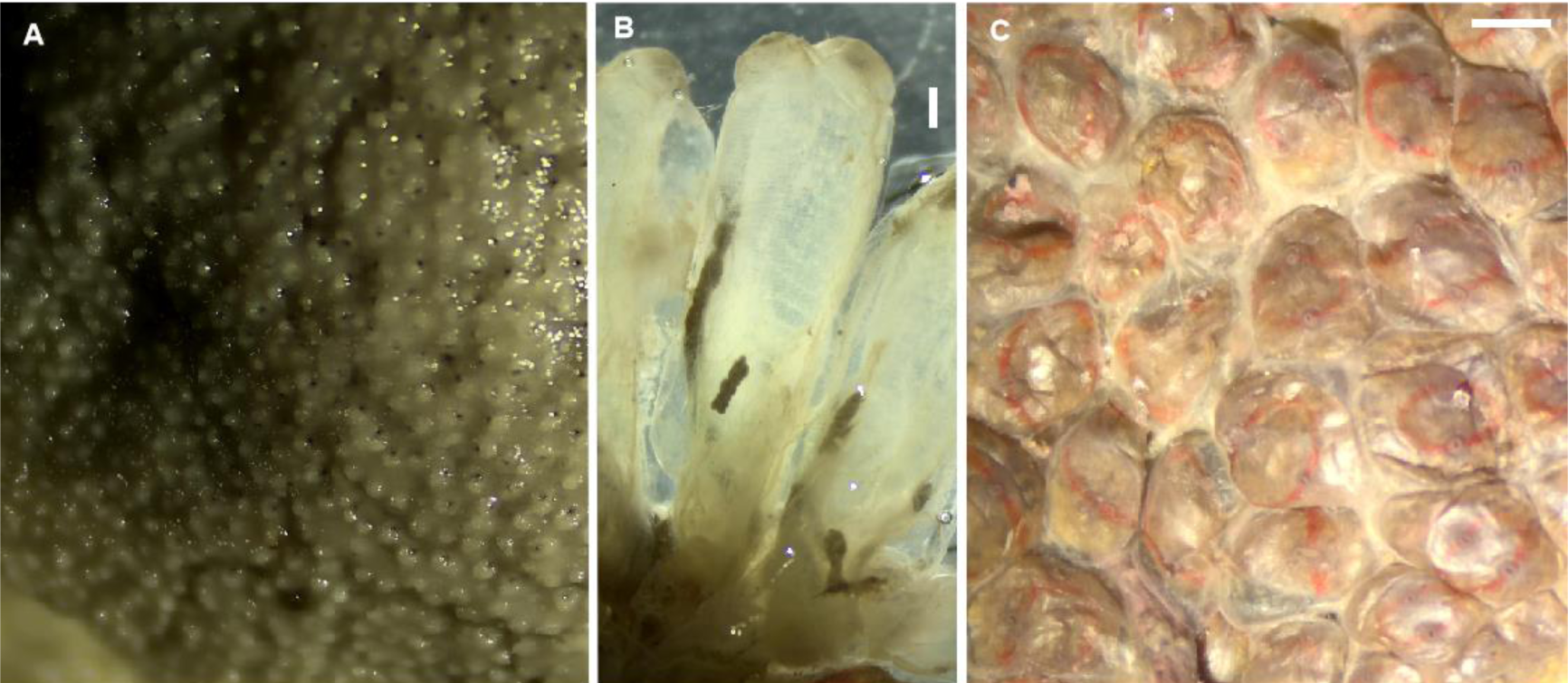
*Didemnum perlucidum* (A), *Ecteinascidia turbinata* (B), *Symplegma rubra* (C). Scale bars: 1mm.

This tunicate species was discovered southwest of Unhocomo Island, where it was observed overgrowing part of a wreck, including a large bivalve. The COI barcode of this colony is 100% identical to samples from Australia, India, the Caribbean, and Turkey. This COI haplotype represents the widely dispersed lineage of *D. perlucidum* (Fig. S1.22), suggesting long-distance dispersal facilitated by boat hulls. Other COI lineages of *D. perlucidum* appear to have more restricted regional distributions (Fig. S1.22). Despite *D. perlucidum* having its type locality inside a marina in the Caribbean (Monniot, 1983), it exhibits higher haplotype diversity in the Indo-Pacific (Fig. S1.22), indicating its non-native status in the Atlantic.

Previously, the cosmopolitan tunicate *D. perlucidum* has been reported only from Senegal (Monniot and Monniot, 1994) and Madeira (Ramalhosa et al. 2021) along the West African coast. This report marks the first detection of the species in Guinea-Bissau.

### Order Phlebobranchia

**Family Perophoridae**

***Ecteinascidia turbinata* Herdman, 1880**

(Fig. 7 B)

Only a single colony of this ascidian species was detected, near Meio Island, on a natural rock substrate. Its COI barcode is 100% identical to those of samples from the Caribbean and the Mediterranean (Fig. S1.23).

*Ecteinascidia turbinata*, previously reported only within the Atlantic, has been documented in West Africa solely in Senegal and Cape Verde Islands (Monniot and Monniot 1994). This study marks the first report of *E. turbinata* in Guinea-Bissau.

### Order Stolidobranchia

**Family Styelidae**

***Symplegma rubra* Monniot C., 1972**

(Fig. 7 C)

This tunicate species was discovered near João Vieira and Meio Islands in the Bijagós, growing on natural rocky substrates. The DNA barcodes of these colonies match those of *Styela rubra* samples from the Caribbean (Fig. S1.24).

*Symplegma rubra* is an Indo-Pacific tropical species reported for the first time in the East Atlantic.

## Conclusions

We documented the first reports of 28 marine invertebrate species in the Bijagós Archipelago. This includes three soft-coral species (considering two putative cryptic species of *Carijoa*), two hard-coral species, fifteen hydroid species (considering two cryptic species under the name *Obelia* aff. *dichotoma*), five bryozoan species, two barnacle species, and three tunicate species. Notable first records for Guinea-Bissau include the corals *Carijoa* spp., *Stragulum bicolor*, and *Tubastrea* spp.; the hydroids *Pennaria* (cf.) *disticha*, *Obelia* aff. *bidentata*, *Obelia* aff. *geniculata*, *Nemalecium lighti*, *Halopteris* cf. *alternata*, *Plumularia* aff. *floridana*, and *Dynamena quadridentata*; the bryozoans of the genera *Schizoporella*, *Bugula*, and *Amathia*; the barnacles *Amphibalanus reticulatus* and *Balanus trigonus*; and the tunicates *Didemnum perlucidum*, *Ecteinascidia turbinata*, and *Symplegma rubra*. Additionally, we provided the first confirmed reports in the East Atlantic for *S. bicolor*, *T. tagusensis*, *N. lighti*, *A. alternata*, *A. distans*, and *S. rubra*. A hydroid species found in the Bijagós, with strong morphological and genetic affinities to *Dyphasia digitalis*, likely represents an undescribed species. These findings highlight a significant deficit in the knowledge of marine biodiversity in West Africa and the biogeography of species flagged as invasive, exotic, non-indigenous, or alien.

Molecular phylogenetic analyses using available DNA barcodes revealed multiple instances of cryptic species diversity within nominal species such as *Carijoa riisei*, *Pennaria disticha*, *Macrorhynchia philippina*, *Obelia bidentata*, *O. dichotoma*, *O. geniculata*, *Nemalecium lighti*, *Halopteris alternata*, *Plumularia floridana*, *P. setacea*, *Dynamena disticha*, *Bugula neritina*, and *Amphibalanus reticulatus*. Recognizing cryptic biodiversity is crucial for accurately understanding or refuting the existence, degree, and origin of potential marine invasions. For example, our analyses suggest that hydroids of the nominal genus *Plumularia* and the nominal species *Obelia geniculata* and *Dynamena disticha* represent complexes of cryptic species with restricted biogeographical distributions, unable to raft long distances on artificial substrates. Consequently, these taxa cannot be flagged as invasive, exotic, or non-indigenous.

Conversely, some nominal species exhibiting cryptic diversity have cryptic species with large geographical distributions due to human influence, as well as cryptic species with restricted ranges unable to cope with drastic abiotic changes for long-distance dispersal. This was the case for hydroids such as *Pennaria disticha*, *Macrorhynchia philippina*, *Obelia bidentata*, *O. dichotoma*, *Halopteris alternata*, the bryozoan *Bugula neritina*, and the barnacle *Amphibalanus reticulatus*. At lower taxonomic levels, we observed nominal species with genetic lineages more widely distributed than others within the same species: the hydroids *Pennaria disticha*, *Macrorhynchia philippina*, *Obelia dichotoma*, *Nemalecium lighti*, and *Halopteris alternata*; the bryozoans *Amathia verticillata*, *Schizoporella errata*, and *Bugula neritina*; the barnacle *Amphibalanus reticulatus*; and the tunicate *Didemnum perlucidum*. Among these, it is notable that cryptic species of the bryozoan *Bugula neritina* and the barnacle *Amphibalanus reticulatus* are widely distributed across warm to cold marine waters. These phylogeographic observations warrant future comparative genomic analyses to reveal adaptive genetic traits that enable or limit the success of some evolutionary units in achieving long-distance dispersal across diverse environmental settings.

The DNA barcoding and morphology-based identification of marine invertebrate species in West Africa are still in the early stages. This study includes the invertebrate taxa we collected from the Bijagós that are more easily assigned taxonomically by DNA barcoding because these species already had determined DNA barcodes elsewhere. We collected and DNA barcoded an even greater diversity of marine invertebrate species during this expedition to the Bijagós (in 2023) than presented here, requiring more detailed taxonomic work (e.g., Moura et al. *in press*; Wirtz et al. *submitted*). This work indicates that marine biodiversity has been grossly overlooked in West African countries, including species flagged as invasive, exotic, or non-indigenous.

We show several examples of great phylogeographic affinities between the warm waters of the Western Atlantic (Brazil, the Gulf of Mexico, and the Caribbean) and West Africa for many of the taxa presented. This is influenced by Atlantic equatorial currents affecting both natural and human-mediated dispersal of marine species, as boat traffic tends to follow these major maritime currents. Thus, we suspect a considerable underestimation of accidental anthropogenic trade of exotic marine species between the tropical and subtropical waters of West Africa and the Atlantic coasts of the Americas, occasionally including the Mediterranean (e.g., *Pennaria* (cf.) *disticha*, *Didemnum perlucidum*). Many exotic species in West Africa were likely introduced much earlier than supposed (e.g., *Nemalecium lighti*). Most alien species in the Tropical East Atlantic originate from the Indo-Pacific, introduced by boats either directly from the SW Indian Ocean or via the Tropical West Atlantic.

A major shipping route connecting southwest Europe and the Indian Ocean via South Africa, which passes only 100 miles west of the Bijagós (cf. https://www.marinetraffic.com/), likely facilitated the direct introduction of many marine exotic species from the Indo-Pacific to West Africa. Additionally, a significant Amphi-Atlantic shipping route connecting the Caribbean and Brazil with the Tropical East Atlantic, roughly between the Cape Verde Islands and Guinea (near the Bijagós; cf. https://www.marinetraffic.com/), likely accounts for most of the unwanted trade of exotic species between the Tropical East and West Atlantic.

Ship traffic within the Bijagós Archipelago is relatively scarce, making direct introductions of exotic species by boats uncommon. Although there might have been instances, especially with recent operations by some Asian fishing fleets near the archipelago, marine exotic species introductions in the Bijagós are more likely due to natural dispersion from other West African marine areas. This suggests that West African coasts should be more heavily infested with exotic species than previously thought.

Among the exotic species identified in the Bijagós, those with the highest potential to disrupt the natural richness of marine biodiversity and affect local fisheries are the bryozoans of the genus *Amathia* (i.e., *A. distans*, *A. verticillata*, and *A. alternata*) and the sun-corals (genus *Tubastraea*). While eradication of these taxa seems impossible, local initiatives to remove *Amathia* bryozoans at low tide may help contain the spread of these invasive species and preserve ecosystem services. Regular monitoring, early detection, and rapid eradication of marine invasions are necessary, not only in Africa but worldwide. It is also urgent to limit the deployment of artificial substrates in the seas and implement more effective measures to prevent the transport of unwanted marine species by boats.

## Data availability

The DNA barcodes determined for the marine invertebrates reported from the Bijagós Archipelago (Guinea-Bissau, West Africa) are available in GenBank under the following accession numbers: PP717971-98, PP718001-12, PP728263-81, PP799165, and PP808917. Additional data can be provided upon request.

## Acknowledgements

We are grateful to the IBAP and INIPO staff in Guinea-Bissau for their invaluable assistance during the expedition. Special thanks to Maria Ana Dionisio, Xavier Turon, and Leandro Vieira for their taxonomic confirmations of selected barnacles, ascidians, and bryozoans. CJM acknowledges the sporadic assistance of various undergraduate students with sorting, photography of samples, and molecular lab work, with special mention to Ema Azevedo, Ana Brito, and Manuel Melo. The fieldwork and exportation of samples were conducted under permit declarations issued by IBAP and INIPO in May 2023.

## Author contributions

Expedition Participation and Organization: CJM, PW, FTN, CB, ES. Sample Collection: CJM, PW. DNA Barcoding: CJM. Article Writing: CJM. Funding Acquisition: ES. Article Revision and Approval: All authors.

