## Supplementary figures and images for "Hotspot of Exotic Benthic Marine Invertebrates Discovered in the Tropical East Atlantic: DNA Barcoding Insights from the Bijagós Archipelago, Guinea-Bissau"

### Fig. S1.1

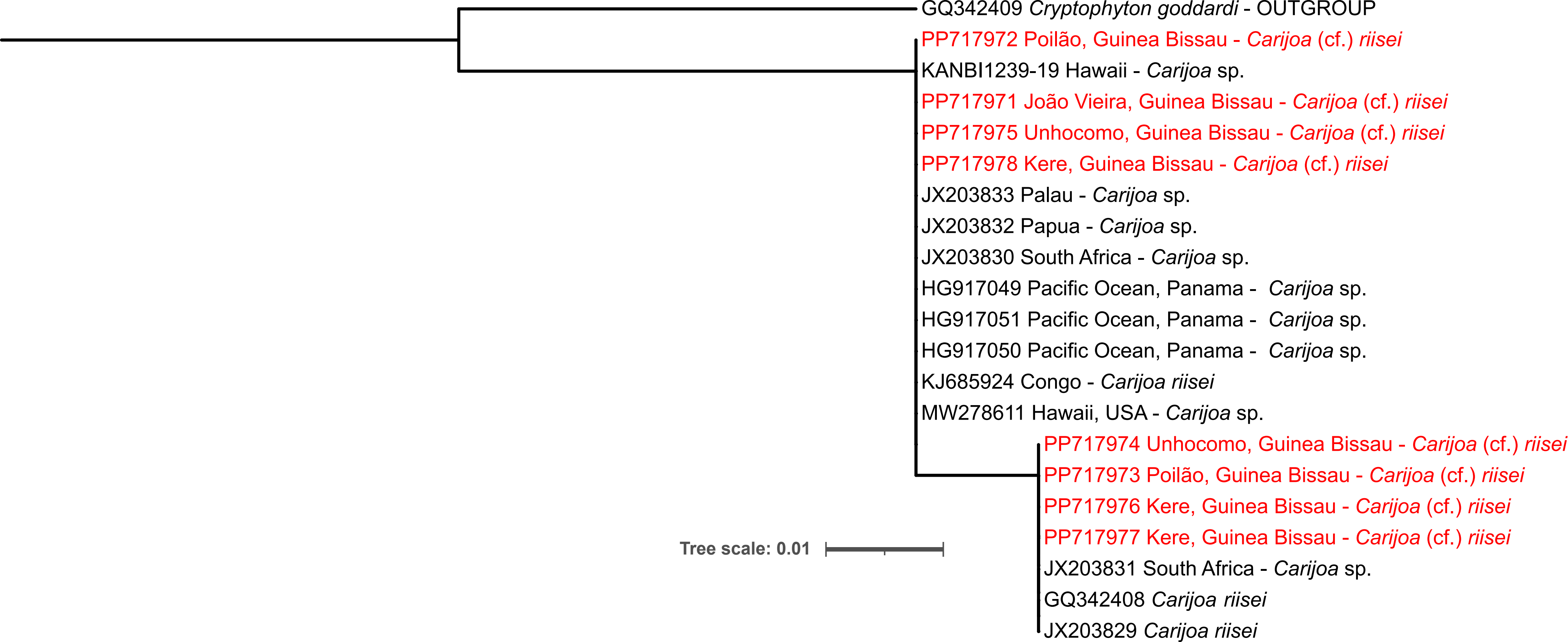

### Fig. S1.2

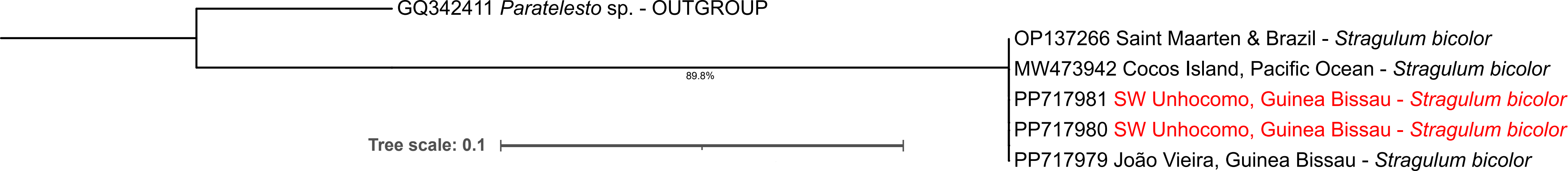

### Fig. S1.3

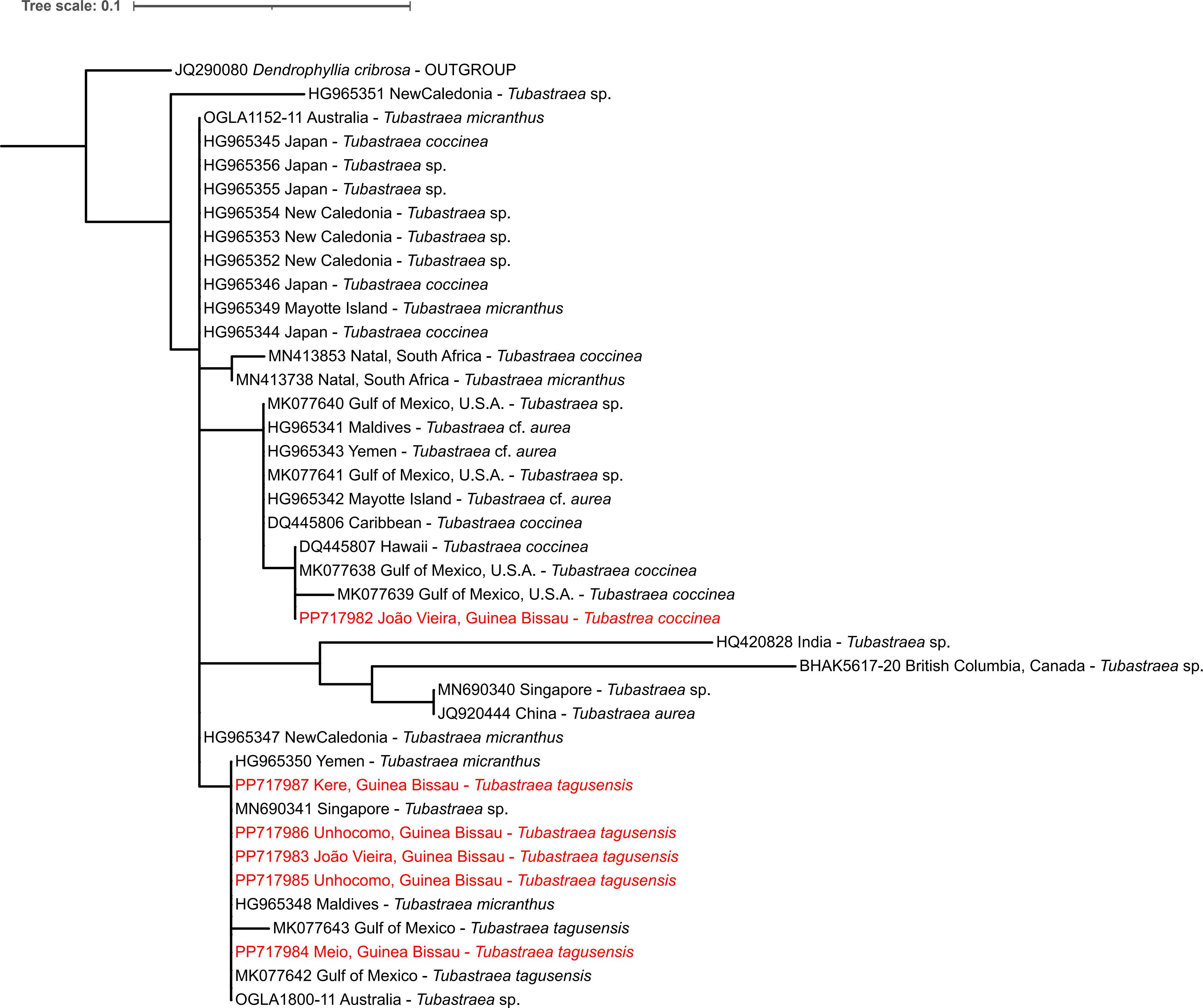

### Fig. S1.4

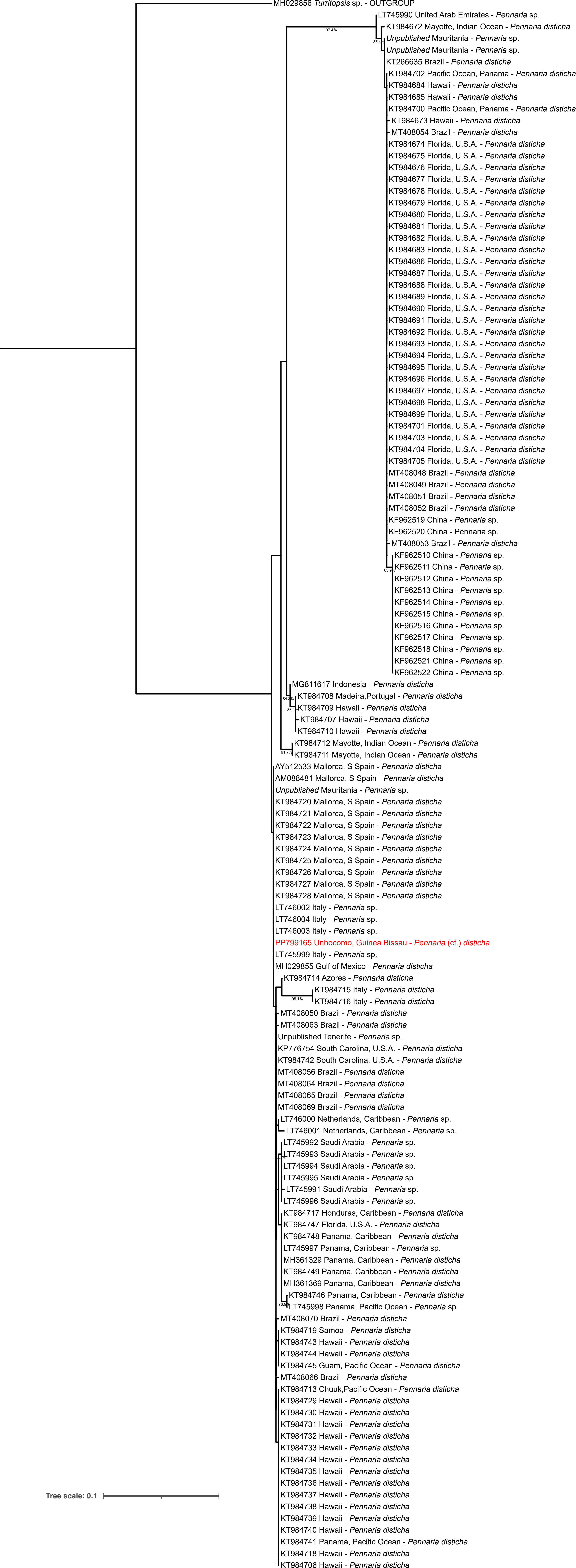

### Fig. S1.5

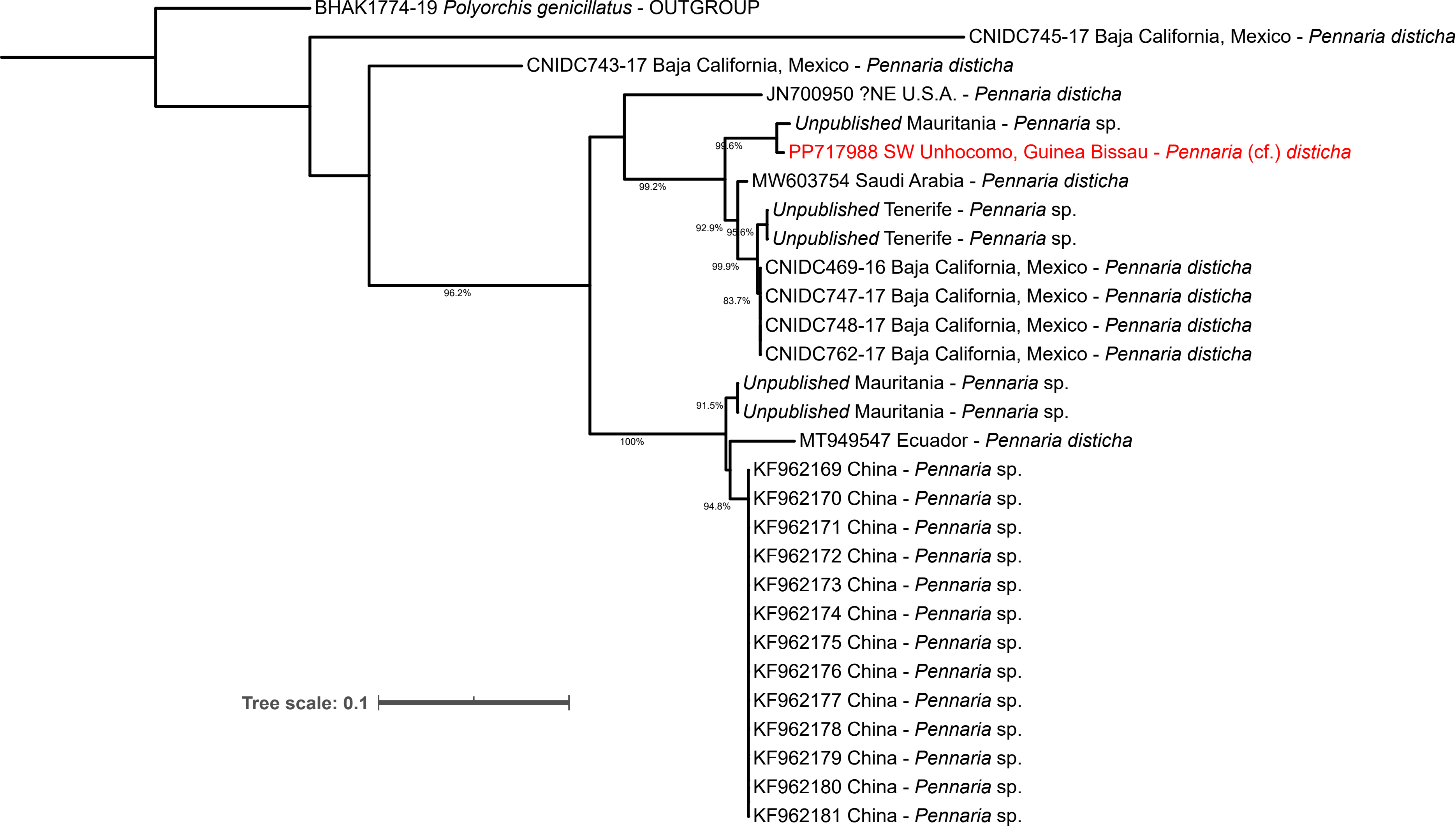

### Fig. S1.6

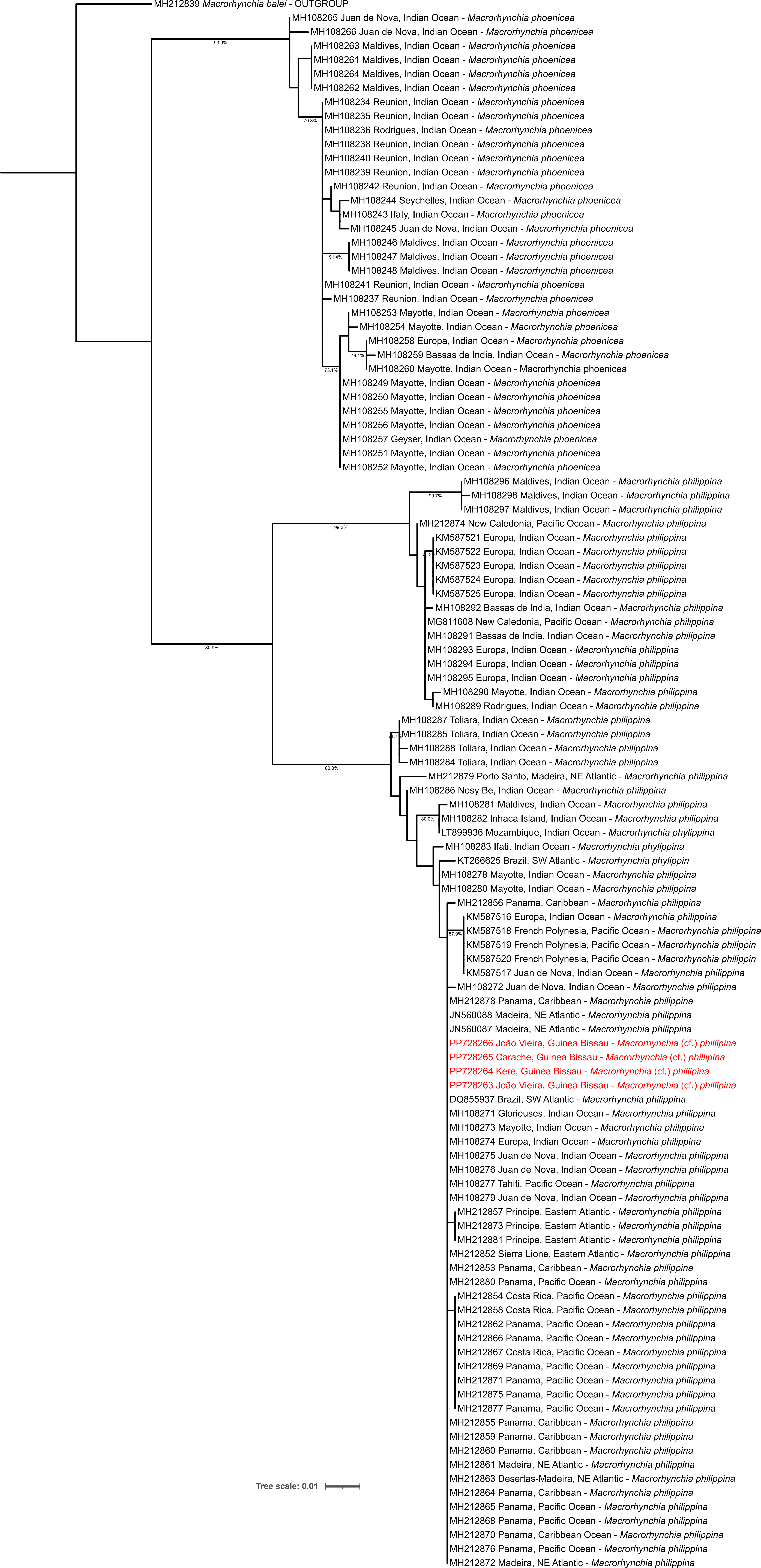

### Fig. S1.7

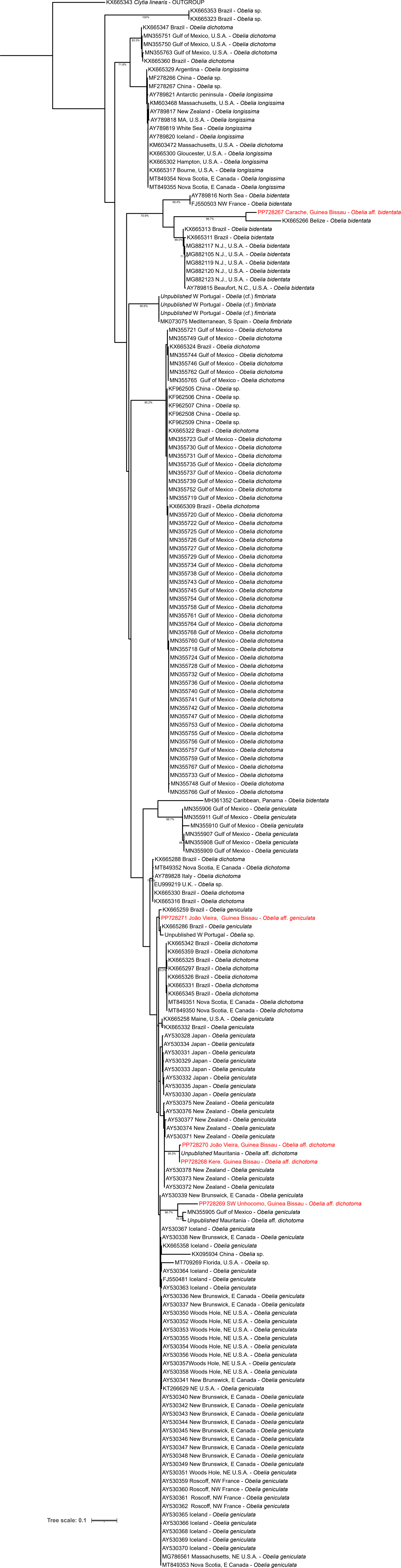

### Fig. S1.8

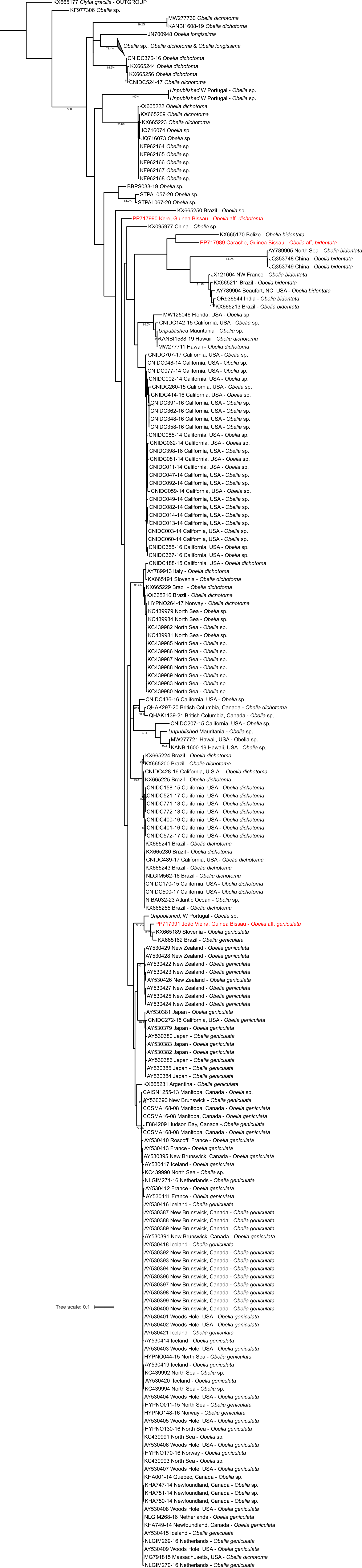

### Fig. S1.9

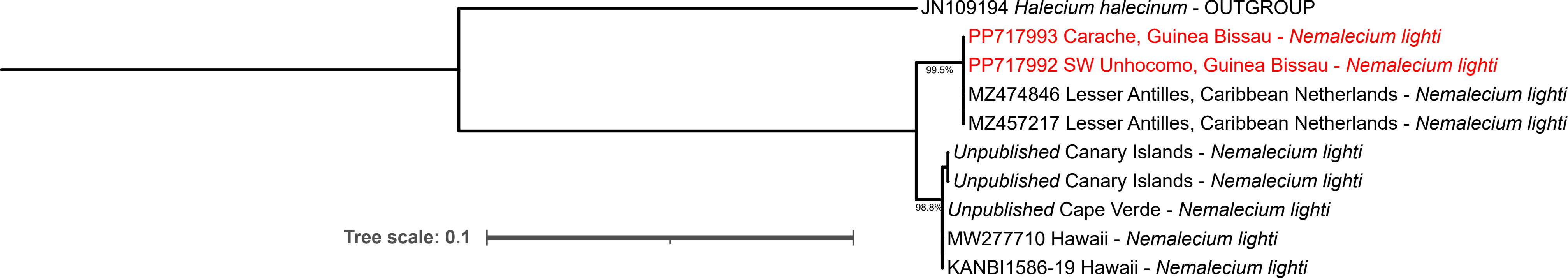

### Fig. S1.10

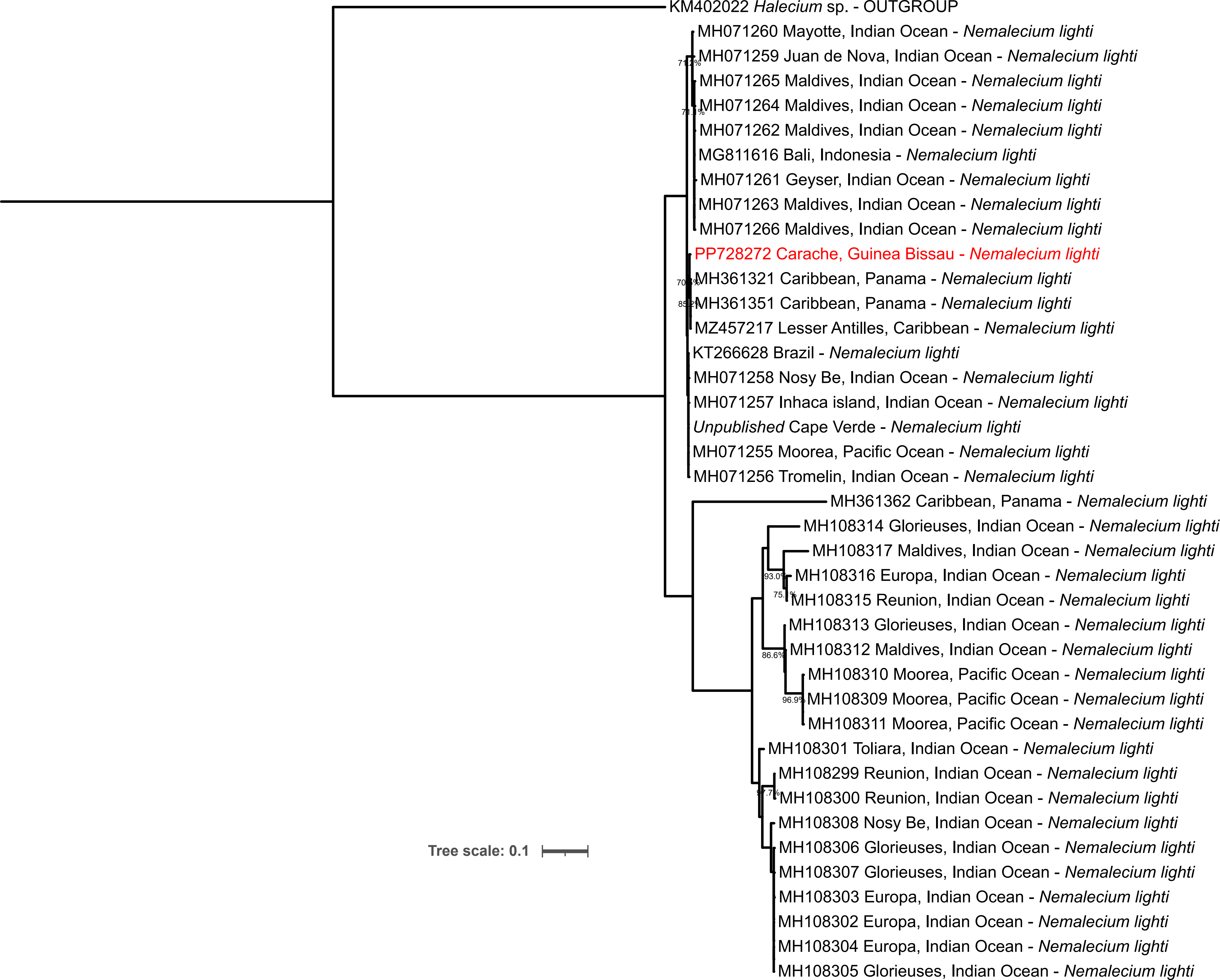

### Fig. S1.11

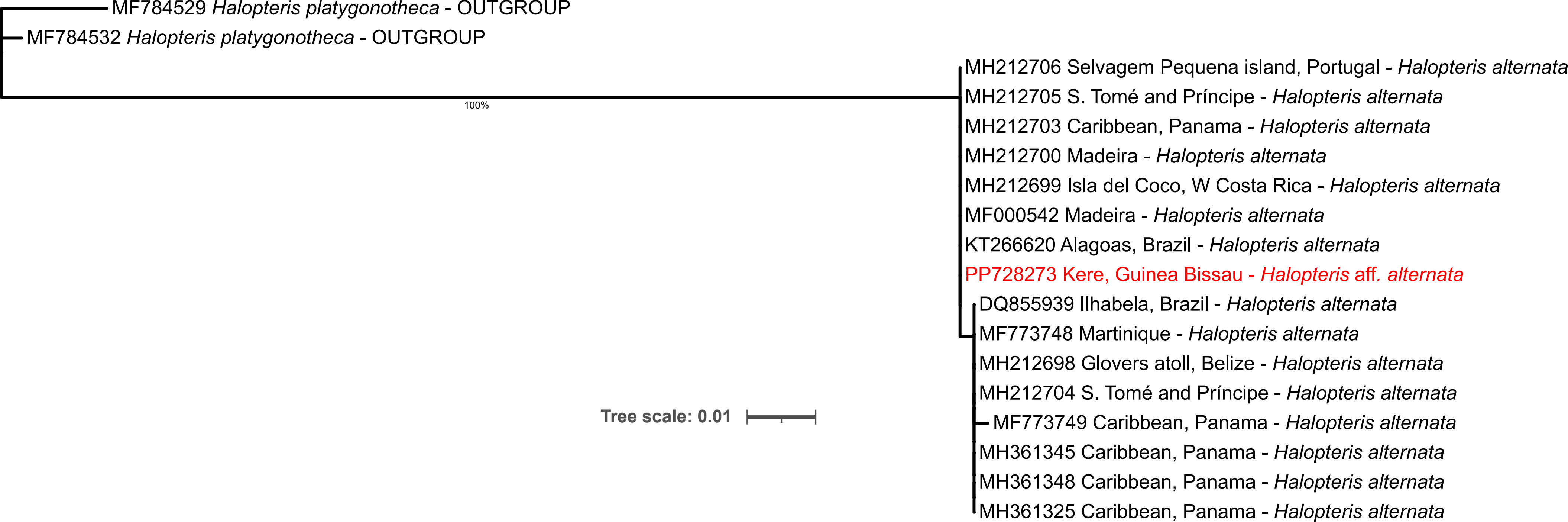

### Fig. S1.12

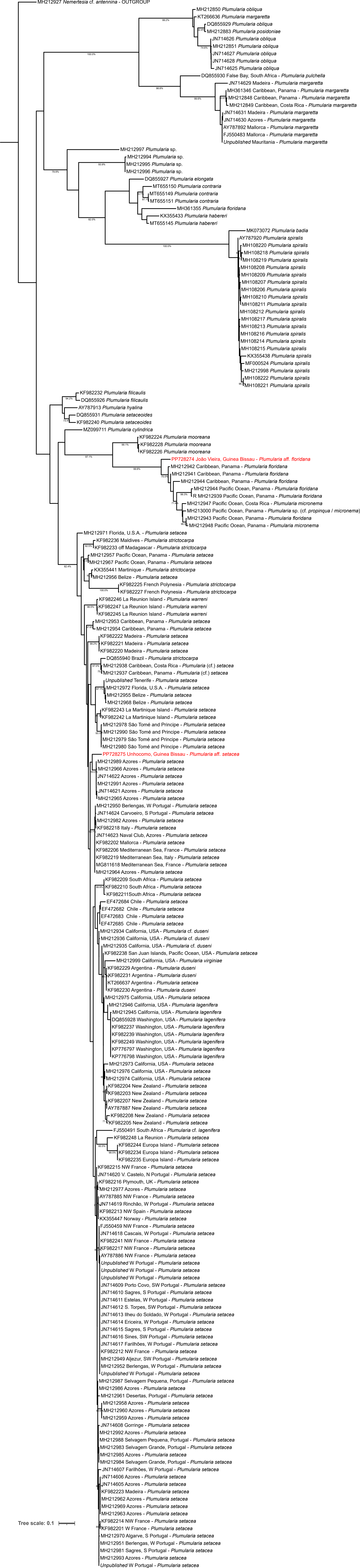

### Fig. S1.13

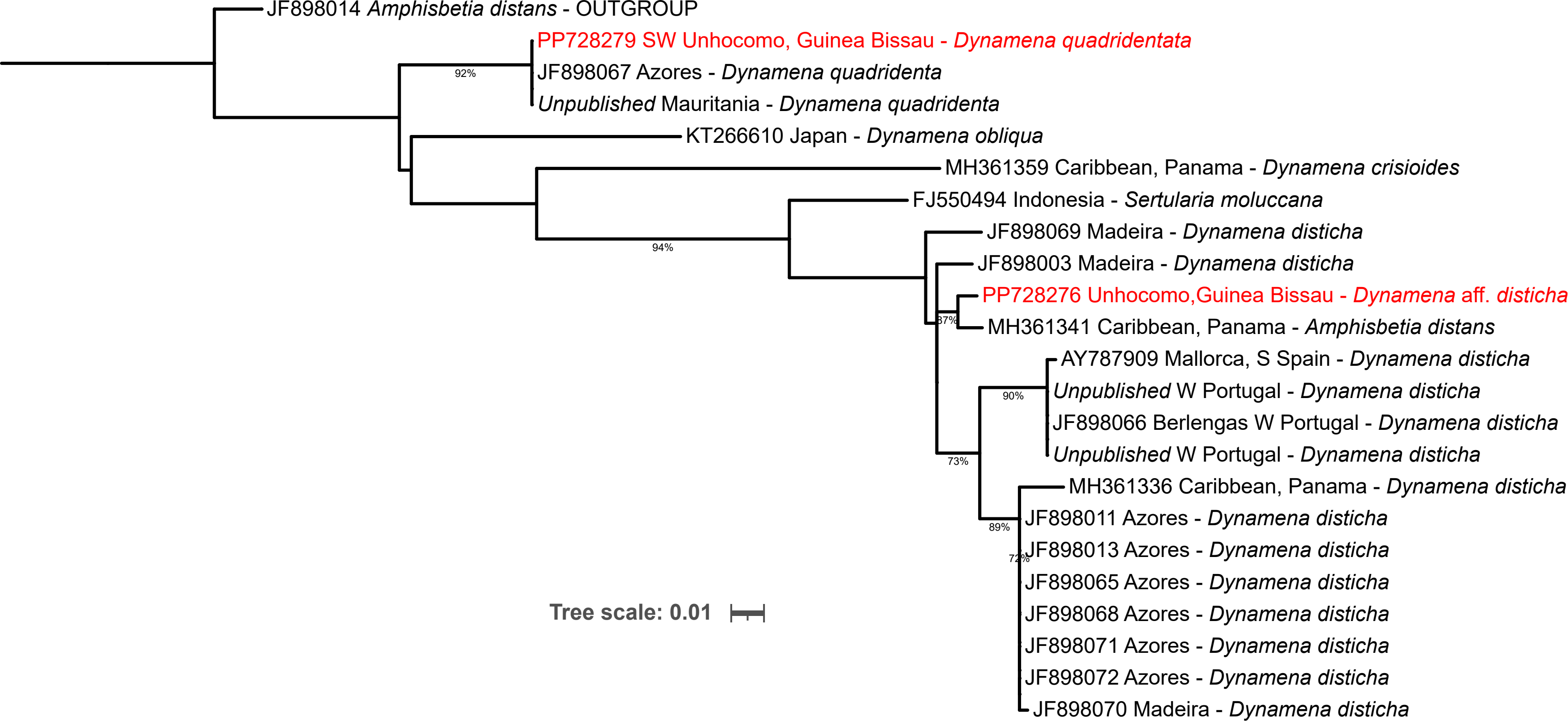

### Fig. S1.14

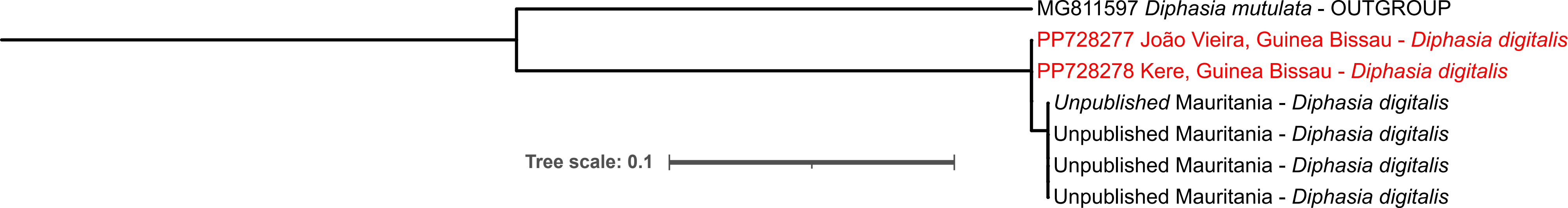

### Fig. S1.15

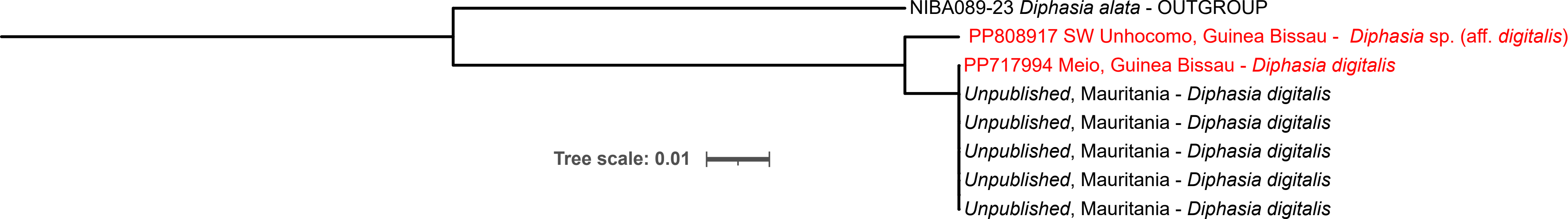

### Fig. S1.16

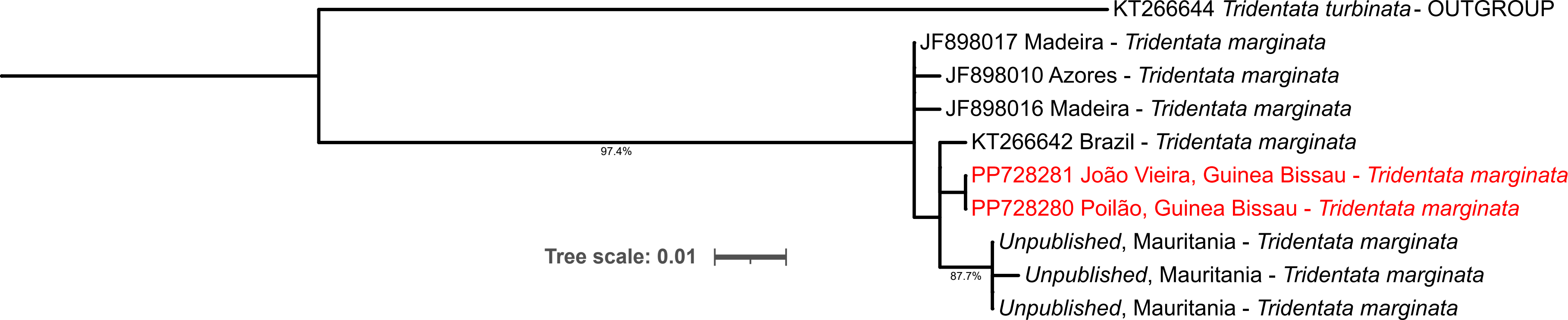

### Fig. S1.17

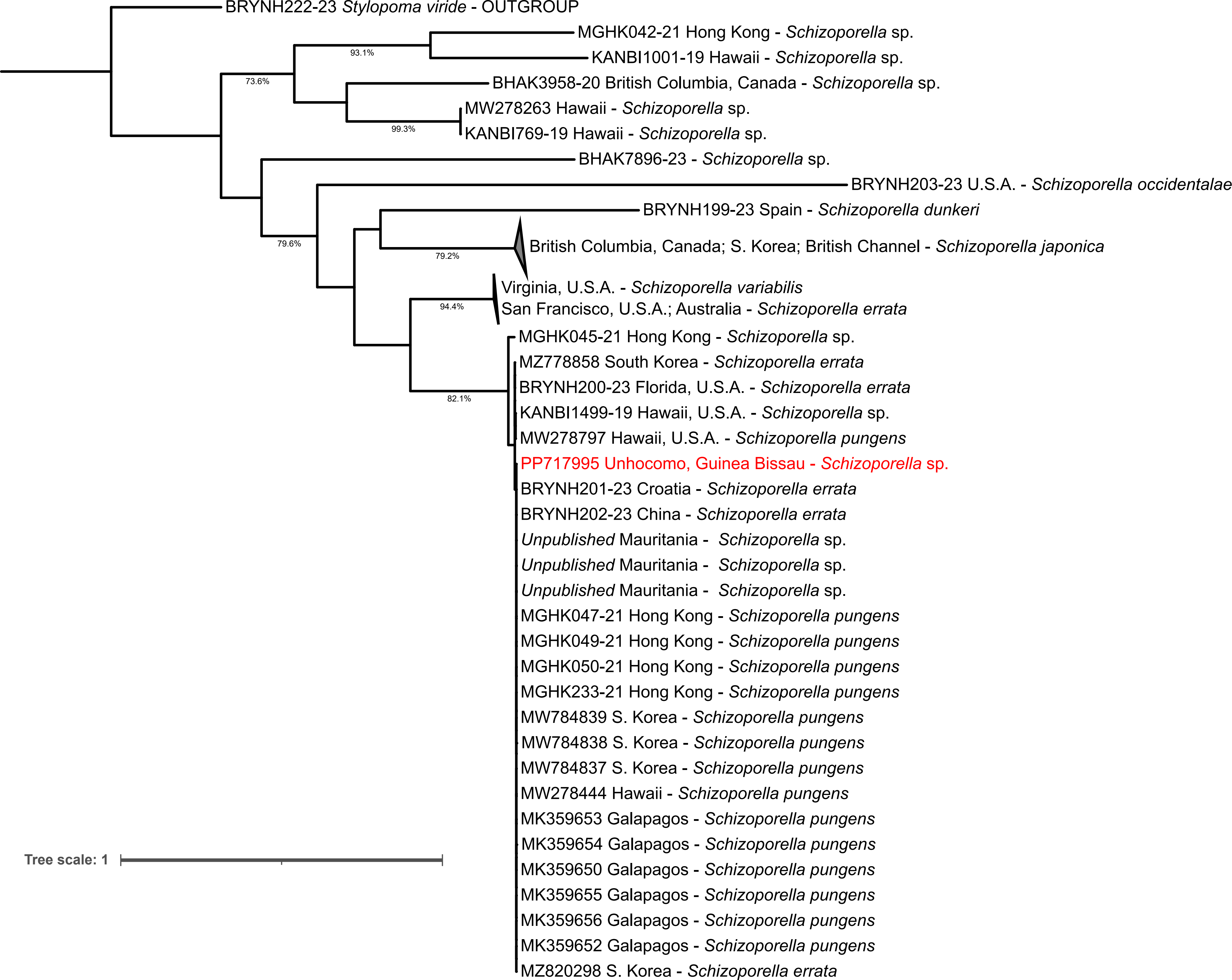

### Fig. S1.18

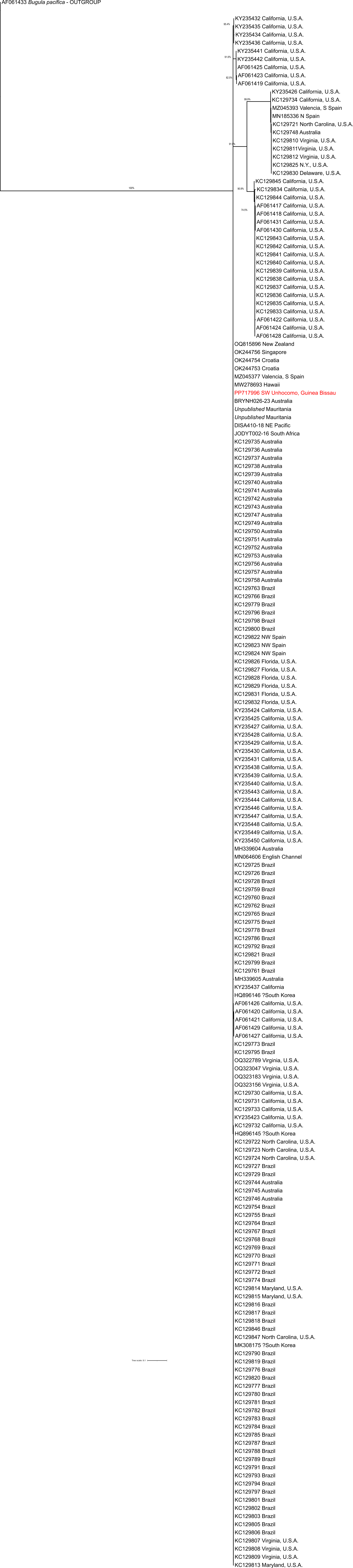

### Fig. S1.19

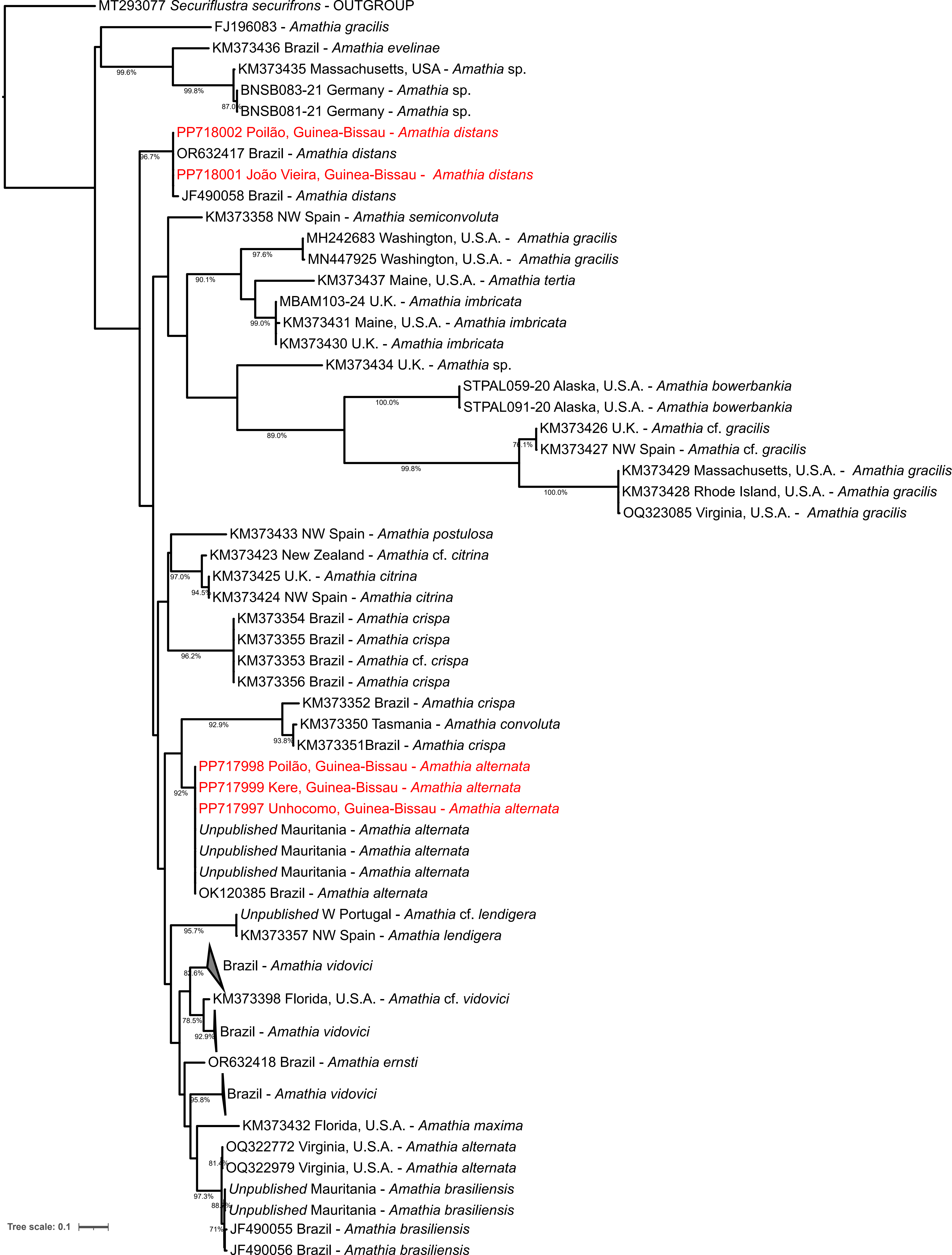

### Fig. S1.20

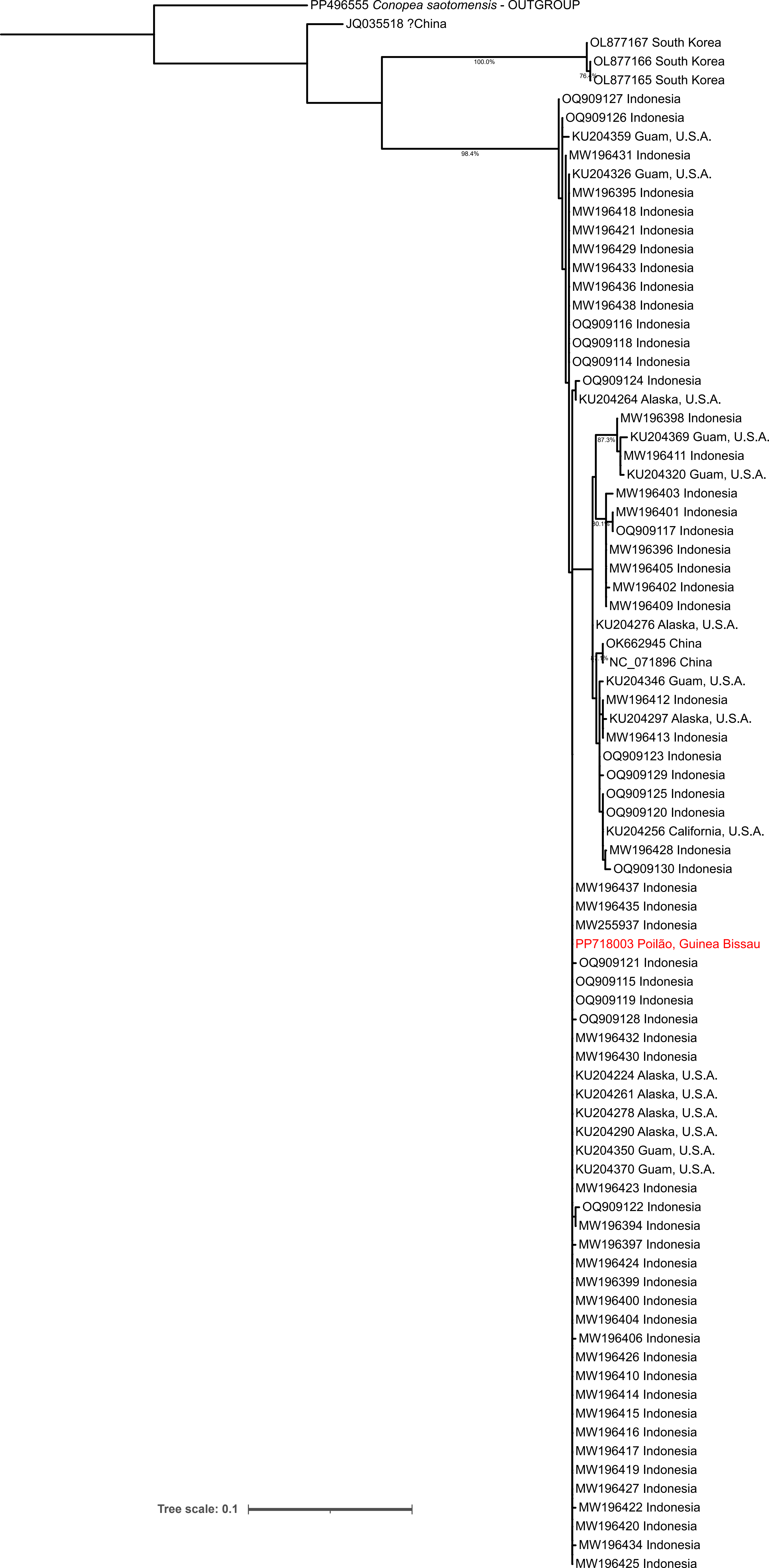

### Fig. S1.21

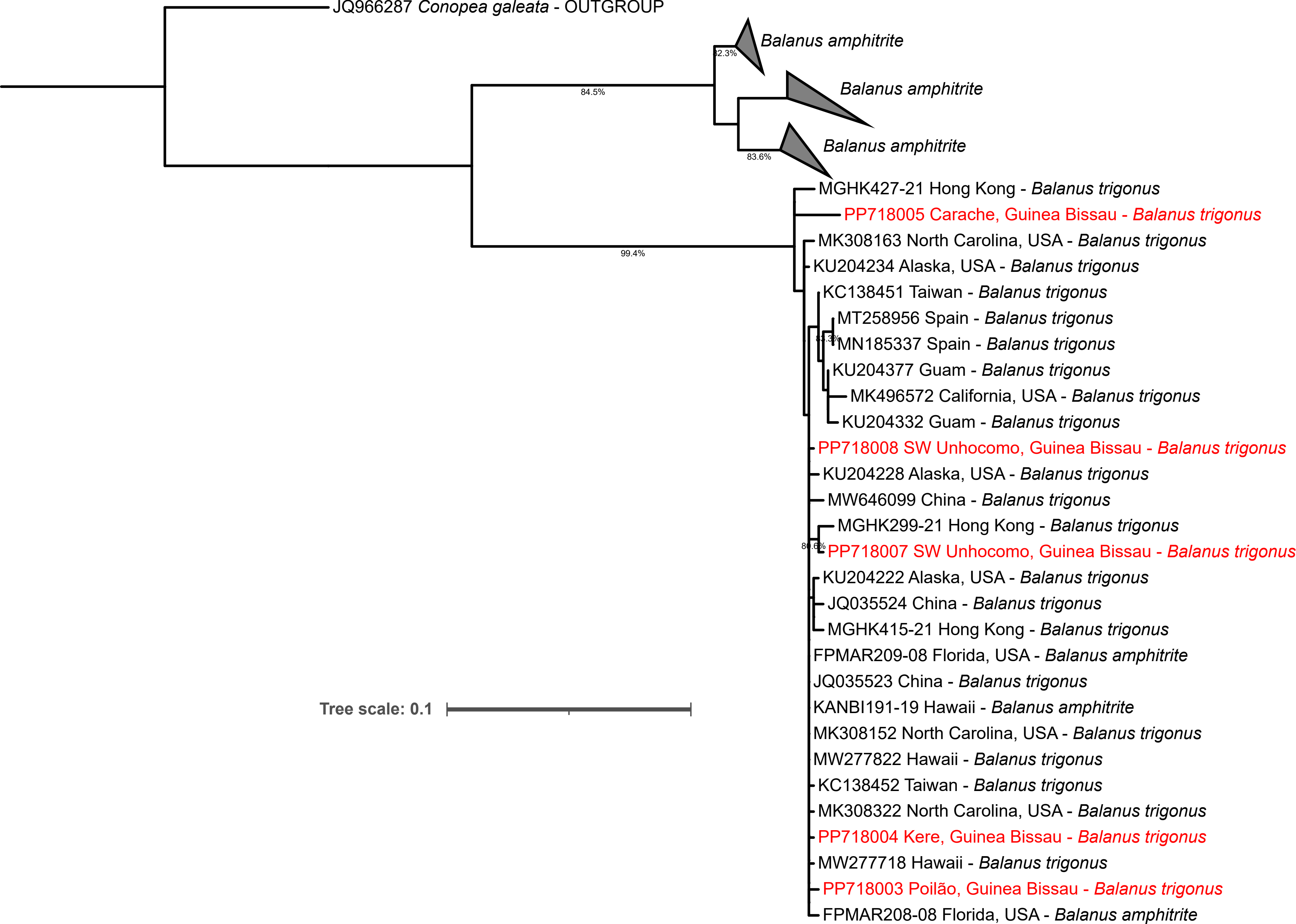

### Fig. S1.22

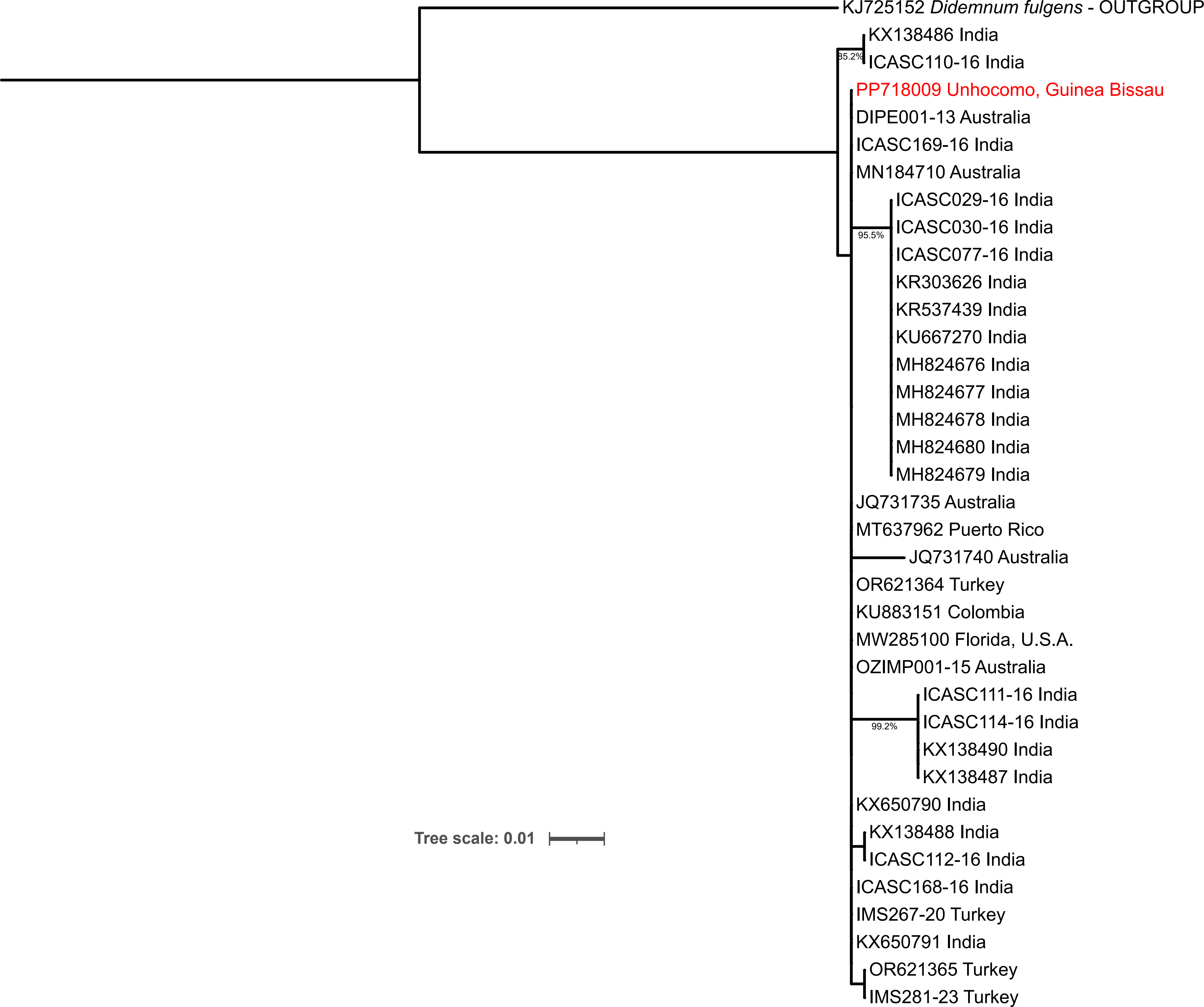

### Fig. S1.23

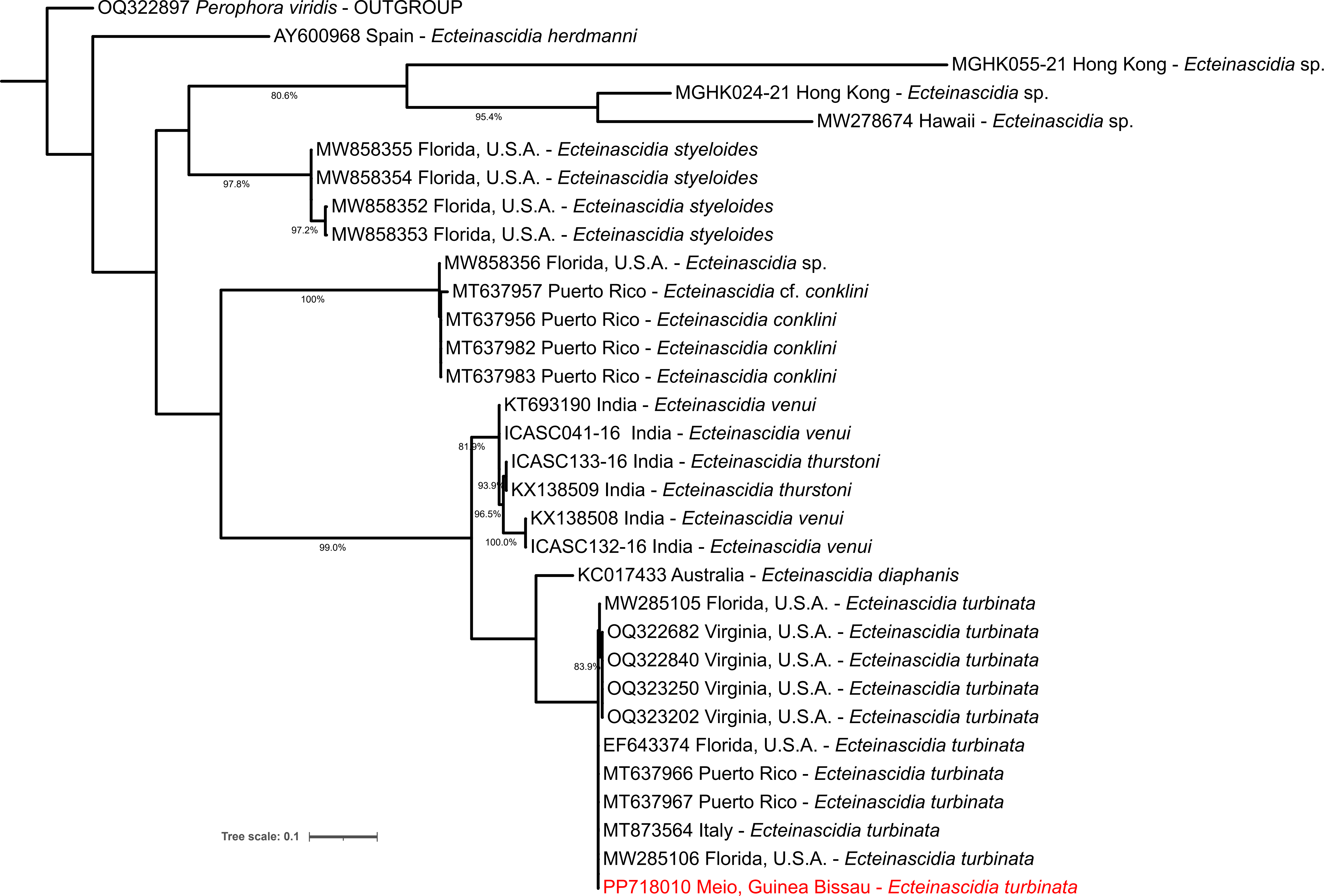

### Fig. S1.24

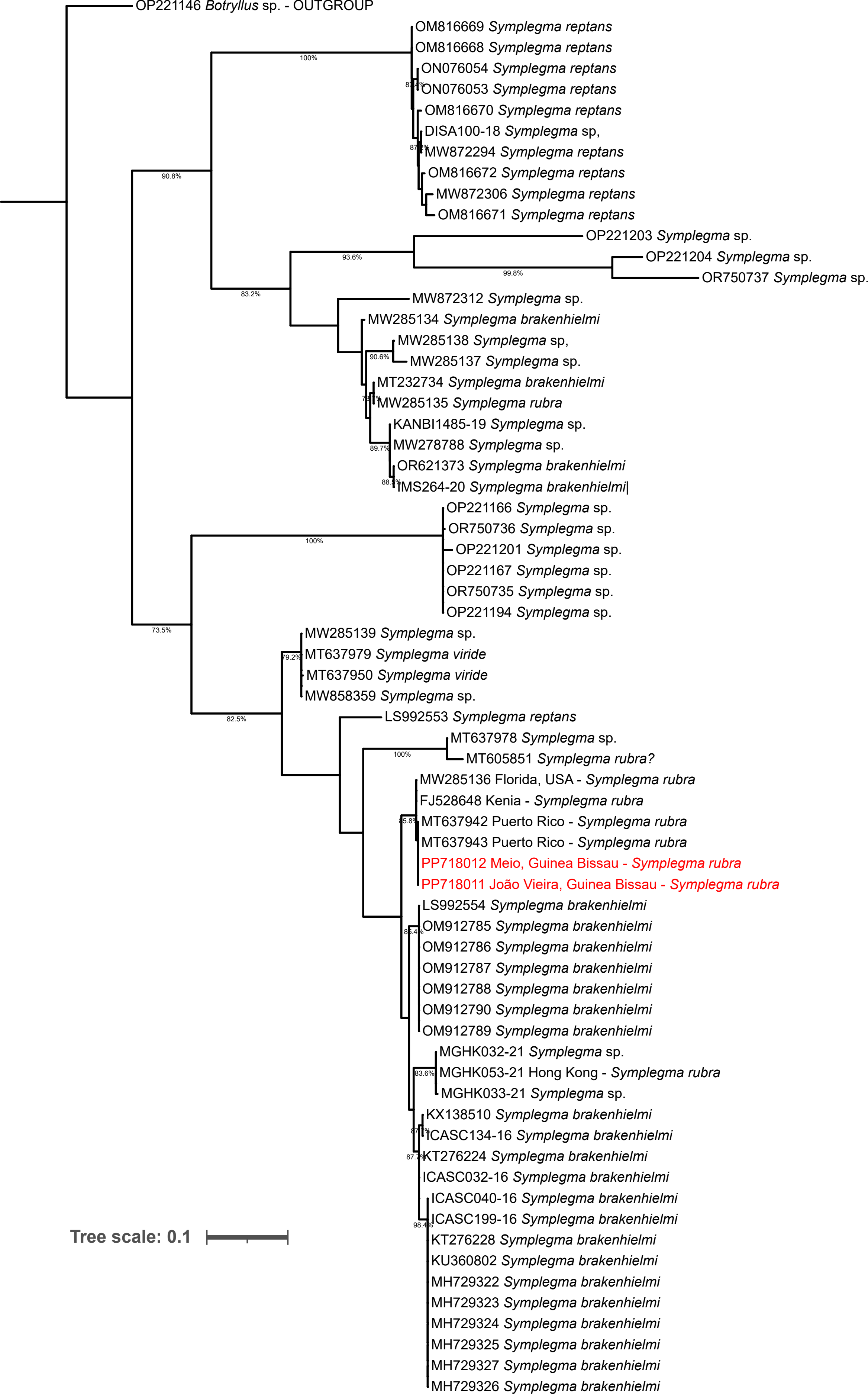
